# Inactivation of tissue-nonspecific alkaline phosphatase reduces phosphatidylcholine amounts in the brain of infant mice

**DOI:** 10.64898/2026.11.12.766160

**Authors:** Lionel G Nowak, Marie Gleizes, Stéphane Balayssac, Océane Delos, Étienne Mornet, Véronique Gilard, Justine Bertrand-Michel, Caroline Fonta

**Author notes:** Corresponding author: Lionel G Nowak; CerCo lab, UMR 5549, Pavillon Baudot, CHU Purpan, BP 25202, 31052 Toulouse Cedex, France.

## Abstract

Hypophosphatasia is a rare disease caused by dysfunctions of tissue-nonspecific alkaline phosphatase (TNAP). TNAP is expressed in numerous tissues, notably the brain, and patients with hypophosphata - sia exhibit neurological symptoms, including epilepsy in the most severe forms of the disease. TNAP knockout (KO) mouse models have provided a wealth of information regarding TNAP functions. Specifically, metabolomic studies focusing on the aqueous fraction of the brain of TNAP KO mice identified numerous molecules whose amounts were altered as a direct or indirect consequence of TNAP inactivation. Among these molecules, several are involved either directly (e.g., vitamin B6) or in - directly (e.g., N-acetylaspartate) in lipid metabolism. In this study, we therefore extended these investi- gations by examining the consequences of TNAP inactivation on the brain lipidome. We used a targeted lipidomic approach, examining four main categories of lipids: fatty acids, neutral lipids, sphin golipids and glycerophospholipids, subdivided into 16 lipid classes and amounting to 296 lipid species. Eleven lipid species showed significantly reduced amounts in the brain of TNAP KO mice. All belonged to the phosphatidylcholine class. As phosphatidylcholines are major constituents of brain cell membranes, our results suggest that hypophosphatasia could be associated with alterations in membrane structure and functions, which could contribute to the neurodevelopmental disorders associated with the disease.

## Introduction

Tissue-nonspecific alkaline phosphatase (TNAP) is a ubiquitous enzyme, which is present in most tis sues and organs. TNAP is an ectoenzyme that cleaves phosphoester bonds, releasing a hydroxyl group and inorganic phosphate (Millán 2006a). TNAP shows broad substrate specificity and is thus able to dephosphorylate a range of extracellular molecules (reviewed in Millán 2006b). Natural extracellular substrates of TNAP include pyrophosphate (PPi), phosphocholine, adenine nucleotides and pyridoxal phosphate (PLP, the active form of vitamin B6). PLP cannot cross cell membranes. TNAP is required for the dephosphorylation of extracellular PLP to pyridoxal (PL), which can be transported into cells where it is rephosphorylated to PLP (reviewed in Coburn 2015). Intracellular PLP, in turn, is a cofactor for several dozen enzymes in mammalians (e.g., Percudani and Peracchi 2009).

In the brain, TNAP is expressed by neurons, astrocytes and oligodendrocytes (reviewed in Fonta et al. 2015). Its activity is highest during development (Goldstein and Harris 1981), in particular in the neurogenic zones during embryonic development (Narisawa et al. 1994; Langer et al. 2007) as well as on axons undergoing myelination (Fonta et al. 2005). However, TNAP activity persists at relatively high level in adulthood (e.g., Friede 1966; Fonta et al. 2004; Langer et al. 2008; Négyessy et al 2011). In adult, prominent TNAP activity persists at the level of synapses in the neuropile and Ranvier’s node of myelinated axons (Sugimura and Mizutani 1979; Mori and Nagano 1985; Fonta et al. 2004, 2005).

In humans, TNAP dysfunction leads to a rare disease called "hypophosphatasia" (Rathbun 1948; Fraser 1957; Whyte 2010; Mornet et al. 2011; Salles 2020). Hypophosphatasia is characterized by diminished alkaline phosphatase activity (Rathbun 1948), which results in elevated urinary and/or serum levels of PLP (Whyte et al. 1985), PPi (Russell 1965) and phosphoethanolamine (PEA) (Fraser et al. 1955). The main medical sign of hypophosphatasia is a defective mineralization of the bone, whose severity depends on the degree of TNAP dysfunction (Fraser 1957; Whyte 2010; Mornet et al. 2011; Salles 2020). Besides, hypophosphatasia is often accompanied by neurological symptoms (Colazo et al. 2019) and in its most severe forms, hypophosphatasia is characterized by the presence of pyridoxine responsive epileptic seizures (e.g., Rathbun 1948; Sia et al. 1975; Baumgartner-Sigl et al. 2007; Taketani et al. 2014). Patients with severe perinatal hypophosphatasia also exhibit abnormalities in global brain structure (Demirbilek et al. 2012; Fukazawa et al. 2018) and may present with multicystic encephalopathy in both white and gray matter (Hofmann et al. 2013; de Roo et al. 2014). Until recent advances in enzyme replacement therapy (Whyte et al. 2016; Bowden and Foster 2019), severe hypophosphatasia was usually fatal.

In the mouse, TNAP is encoded by the *Akp2* gene. The *Akp2* knockout mouse model resumes severe lethal perinatal hypophosphatasia, including bone mineralization defects and pyridoxine responsive epileptic seizures (Waymire et al. 1995; Narisawa et al. 1997), as well as elevated levels of plasmatic and/or urinary PLP, PEA and PPi (Waymire et al. 1995; Fedde et al. 1999). Histological examination of the nervous system of *Akp2* KO mice revealed thinning of lumbar nerve roots (Narisawa et al. 1997) as well as altered synaptogenesis in the brain and defective myelination in the brain and spinal cord (Hanics et al. 2012).

Analysis of brain extracts of *Akp2*^-/-^ mice initially revealed a drastic reduction in the concentration of GABA, the principal inhibitory neurotransmitter (Waymire et al. 1995). In a subsequent untargeted metabolomics study (Cruz et al. 2017) examining the aqueous fraction of brain extracts, we confirmed the decrease in GABA level (-40%) and further disclosed significant alterations in concentration for 7 other metabolites in *Akp2*^-/-^ mice: cystathionine (+290%), adenosine (-86%), 3-methylhistidine (+74%), methionine (+50%), histidine (+20%), N-acetylaspartate (NAA, +17%), and N-acetyl-aspartylglutamate (NAAG, +13%) (see also Phoenix et al. 2025). Since both GABA and adenosine are involved in inhibiting neuronal activity (Phillis et al. 1979; Connors et al. 1988), their decreased amounts offer an explanation for the severity of epilepsy in the severe forms of hypophosphatasia. The change in adenosine level could be attributed to the ectonucleotidase function of TNAP while the changes in the amounts of GABA, cystathionine, histidine, 3-methylhistidine, and methionine could be attributed to the direct or indirect consequences of altered PLP-dependent enzyme activities.

The present study was aimed at exploring the lipidome of *Akp2* KO mice. The high abundance of lipids in the brain makes it the second most lipid-rich organ after adipose tissue (e.g., O’Brien and Sampson 1965a; Yoon et al. 2022). The great diversity of lipid chemical structures gives them a wide range of functions: in addition to their roles in energy metabolism, lipids determine the structure and shape of cellular membranes; lipids are also precursors of second messengers, of signaling molecules and of mediators of inflammation; through their interactions with proteins, they ensure essential membrane dynamics such as vesicle release or neurite growth; lipids also regulate the activity of membrane bound enzymes and transmembrane ion channels (reviewed in: Farooqui et al. 2000a; Lauwers et al. 2016; Harayama and Riezman 2018; Yoon et al. 2022).

Several results from previous studies led us to hypothesize that TNAP dysfunction could also result in alterations in lipid metabolism. First, we expected that sphingolipid levels would be reduced in the brains of TNAP KO mice. Indeed, the first step in the *de novo* synthesis of all sphingolipid species depends on a PLP-dependent enzyme, serine palmitoyltransferase (Braun et al. 1970; Hanada 2003; Bourquin et al. 2011), which catalyzed the condensation of serine and palmitoyl-CoA to produce 3-ketodihydrosphingosine (sphinganine). Since TNAP dysfunction leads to decreased brain PLP concentration (Waymire et al. 1995), we expected that all sphingolipid species would be affected in *Akp2^-/-^*mice.

Second, we expected a decrease in the level of myelin-specific lipids in the brains of TNAP KO mice. Indeed, we previously reported that N-acetylaspartate (NAA) and N-acetyl-aspartylglutamate (NAAG) accumulate in the brains of *Akp2* KO mice (Cruz et al. 2017), even though their metabolism is independent of PLP-dependent enzymes. NAA and NAAG are among the most abundant metabolites of the brain and have been given several putative roles (reviewed in Baslow and Guilfoyle 2013; GrønbækThygesen and Hartmann-Petersen 2024). Among these, it has been proposed that NAA, synthesized by neurons, is taken up by oligodendrocytes – the brain cells responsible for myelin formation –, where it is catabolized into aspartate and acetyl-CoA. Acetyl-CoA would then be incorporated into the fatty acids necessary for the synthesis of myelin lipids (reviewed in Namboodiri et al. 2006). Although questioned (Baslow and Guilfoyle 2013; Grønbæk-Thygesen and Hartmann-Petersen 2024), this hypothesis suggests that accumulation of NAA and NAAG in the brains of TNAP KO mice could be the conse quence of a decrease in lipid production, especially for lipids involved in myelin synthesis. This there fore predicted a decrease in myelin specific lipids in the brains of *Akp2*^−/−^ mice.

Third, we expected that amounts of phosphatidylcholines (PCs), which are among the most abundant brain lipids (Wells and Dittmer 1967; Jain et al. 2014; Fitzner et al. 2020), would be decreased in the brains of *Akp2*^−/−^ mice. This rests on the observation that methionine and cystathionine accumulate in the brains of *Akp2*^−/−^ mice (Cruz et al. 2017; Phoenix et al. 2025). This accumulation indicates profound perturbation of the transsulfuration pathway and of the methionine cycle ( Cruz et al. 2017). The methionine cycle is the main source of methyl groups used for the methylation of a wide range of molecules including proteins and DNA (reviewed in Kloor and Osswald 2004; Loenen 2006). Notably, methylation is required in one of the two PC synthesis pathways (reviewed in: Farooqui et al. 2000a; van der Veen et al. 2017). In this synthesis pathway, PCs are produced after the triple methylation of phosphatidylethanolamines (PEs) by phosphatidylethanolamine methyl transferase (PEMT). This suggests that inactivation of TNAP, by perturbing the methionine cycle, could indirectly impact on PCs synthesis by PEMT.

In order to scrutinize these putative consequences of TNAP dysfunction, we analyzed the lipidome of the brains of *Akp2*^+/+^, *Akp2*^+/-^ and *Akp2*^-/-^ mice. Sixteen lipid classes were examined, within which 130 lipid species were identified out of 296 targeted species. Among these 16 classes, decreases were found only for species of the PC class. It has been shown that decreases in PC levels have multiple consequences such as disruption of the integrity of cell membranes (Li et al. 2006), reduction of neuronal process growth (Posse de Chaves et al. 1995a) and impairment of lipid transport (van der Veen et al. 2017). Our study therefore suggests that hypophosphatasia may have hitherto unrecognized consequences on lipid dependent brain functions.

## Methods

### Animals and sample collection

The present study is based on a subsample of the sample of brains used in the metabolomics study of Cruz et al. (2017). *Akp2* mice (Narisawa et al. 1997) were bred in the CerCo animal facilities in accordance with the guidelines for the care and use of laboratory animals (National Research Council 1996, European Directive 86/609) and the guidelines of the local institutional animal care and use committee.

The study was approved by the regional (Midi-Pyrénées, France) ethics committee (ref. No. MP/06/79/11/12). In our rearing conditions, *Akp2*^−/−^ mice did not survive beyond 8-10 days. The brains were therefore taken from 7-day-old animals. For sample collection, mice were weighed, anesthetized by hypothermia to slow down metabolism, and then decapitated. Brains, including olfactory bulbs, cerebral hemispheres, brainstem and cerebellum, were quickly removed, weighed, immediately frozen in liquid nitrogen and stored at –80°C. Mouse genotypes were specified *a posteriori* using tail samples (for details on genotyping see Cruz et al. 2017). Analysis of the lipidome was made on a total of 15 brains, 5 from wild-type *Akp2*^+/+^, 5 from heterozygous *Akp2*^+/–^, and 5 from homozygous knockout *Akp2*^−/−^ mice.

### Chemicals

Internal standards used for lipidomic analysis were purchased from Avanti Polar Lipids. Solvents were purchased from Fisher Scientific.

### Tissue extraction

The frozen tissues were weighed and the brain extracts prepared according to the procedure described by Beckonert et al. (2007) using a chloroform:methanol:water mixture [2:2:1.425 (v/v/v)]. The whole brain was transferred into an ice-cold glass vial maintained on ice. Ultra-pure cold water (0.425 mL) and cold methanol (2 mL) were added. The suspension was pulverized with an Ultra-Turrax disperser (IKA-Werke, Staufen, Germany) for 3 × 10 s and then sonicated with a sonicator probe (Vibra cellTM, Sonic, Newtown, USA) for 3 × 10 s. Two mL of cold chloroform and 1 mL of ultra-pure cold water were then added and the mixture was vortexed for 15 s and kept on ice for 15 min. After centrifugation (15 min, 1100 g, 5000 rpm, 4°C), the lower chloroformic phase was collected for lipidomics. The solution was then aliquoted into 3-mg-tissue equivalents for each lipidomic analysis. Each aliquot was subsequently evaporated to dryness.

### Lipidomics

#### Neutral lipids analysis

Before analysis, internal standards (6 µg of stigmasterol; 6 µg of cholesteryl ester C17:0; 16 µg of tria cylglycerol C57:0) were added to the extract. One µL of the lipid extract was analyzed by gas chromatography flame ionization detector on a GC Trace 1300 (Thermo Electron) system using an RTX-5 column (5% polysilarylene, 95% polydimethylsiloxane, 5 m × 0.25 mm i.d., 0.25 µm film thickness) (Barrans et al. 1994). Oven temperature was programmed from 190°C to 350°C at a rate of 5°C/min and the carrier gas was hydrogen (5 mL/min). The injector and the detector were at 315°C and 345°C respectively. Peak detection and integration were performed using Chromeleon software (Thermo Scientific).

#### Phospholipid analysis

Phospholipid internal standards (PE 12:0/12:0, 180 ng; PC 13:0/13:0, 20 ng; phosphatidylinositol (PI) 15:0/18:1, 30 ng; phosphatidylserine (PS) 12:0/12:0, 1600 ng) were added to the extracts, dried under nitrogen and reconstituted in 50 µL of MeOH. The extracts were analyzed using an Agilent 1290 ultra performance liquid chromatography (UPLC) system coupled to a G6460 triple quadrupole mass spec trometer (Agilent Technologies). A Kinetex HILIC column (Phenomenex, 50 × 4.6 mm, 2.6 µm) was used for LC separations. The column temperature was set at 40°C. The mobile phase A was acetonitrile and the mobile phase B was 10 mM ammonium formate in water at pH 3.2. For PE and PC, the gradi ent was as follows: from 10% to 30% B in 10 min, 100% B for 10-12 min, and then back to 10% B at 13 min for 2 min prior to the next injection. The flow rate of the mobile phase was 0.3 mL/min and the injection volume was 2 µL. For PI and PS, the gradient was as follows: from 5% to 50% B in 10 min and then back to 5% B at 10 min for 9 min prior to the next injection. The flow rate of mobile phase was 0.8 mL/min and the injection volume was 5 µL. Electrospray ionization was performed in positive mode for PE and PC analysis and in negative mode for PI and PS analysis. Needle voltage was set at 4 kV and -3.5 kV respectively. The analyses were performed in Selected Reaction Monitoring (SRM) detection mode using nitrogen as collision gas. Ion optics and collision energy were optimized for each lipid class. Peak detection, integration and quantitative analysis were performed using MassHunter Quantitative Analysis software (Agilent Technologies).

#### Total fatty acids methyl ester analysis

After the neutral lipids and phospholipids analysis, the extracts were hydrolyzed with KOH 0.5 M in methanol (1 mL) at 55°C for 30 min, and transmethylated in boron trifluoride methanol solution 14% (1 mL) and heptane (1 mL) at 80°C for 1 h. After addition of water (1 mL) to the crude, fatty acids methyl esters (FAMEs) were extracted with heptane (2 mL), evaporated to dryness and reconstituted in ethyl acetate (20 µL). FAMEs (1 µL) were analyzed by gas chromatography (Trace 1300, Thermo Fisher) equipped with a flame ionization detector (Lillington et al. 1981) using Famewax RESTEK fused silica capillary column (30 m × 0.32 mm i.d, 0.25 µm film thickness). Oven temperature was programmed from 100°C to 250°C at a rate of 6°C/min and the carrier gas was hydrogen (1.5 mL/min). The injector and the detector were at 220°C and 230°C respectively. Finally, peak detection and integration were performed using Chromeleon software (Thermo Scientific). The internal standard was tria cylglycerol (TAG) 19:0 (4 µg).

#### Sphingolipid analysis

Sphingolipids internal standards (64 ng of sphingomyelin 30:1;O2; 64 ng of glucosylceramide 18:1;O2/12:0; 48 ng of ceramide 18:1;O2/12:0; 80 ng of lactosylceramide 18:1;O2/12:0; 20 ng of dihydroceramide d18:0/12:0; 280 ng of sphingosine 17:1 and sphinganine 17:0) were added to the extracts. The extracts were solubilized in 250 µL of NaOH 0.5 M/MeOH and 250 µL of CHCl_3_ and heated at 37°C for 2 hours. Then, 250 µL of HCl 0.5M/MeOH, 430 µL water, 850 µL of CHCl _3_ and 500 µL of CHCl_3_/MeOH (2:1 v/v) were added. The organic phase was washed with 1 mL of CHCl_3_/MeOH/H_2_O (3:48:47 v/v/v). The organic phase was finally evaporated under nitrogen and dissolved in 20 µL of MeOH. All acquisitions were performed using an Orbitrap Exploris 240 mass spectrometer (Thermo Scientific) coupled with the Waters Acquity UPLC Class I system. An Acquity UPLC CSH C18 column (Waters, 1.0 × 100 mm) was used for LC separation. The mobile phase consisted of 0.1% formic acid, 1 mM ammonium formate in water:MeOH 70:30 (A), as well as in MeOH:IPA 70:30 (B) at a flow rate of 0.1 mL/min and a column temperature at 50°C. Linear gradient solvent composition was used for high-performance liquid chromatography starting with 50% B and raised to 99% B in 8 min, which was maintained at 99% B for 3 min and then decreased back to 50% B in 0.5 min, 4.5 min equilibration time was given. Five µL of the samples were injected into LC. The source settings used were the following: spray voltage +3.4 kV, capillary temperature 320°C, sheath gas 25 and auxiliary gas 5 (arbitrary units). For full mass, the resolution was set at 120,000 full width at half maximum height (FWHM) within the scan range from m/z 200–1400, and automatic gain control (AGC) target, 10^6^. For the stage of data-dependent MS/MS, fragmentation was realized on ions up to 10^6^ under stepped Collisional Energy (10, 15, 20, 30 eV). The settings used were the following: resolution of 30,000 FWHM, AGC target of 10^6^, and isolation window of 1.6 m/z. Peak detection, integration and quantitative analysis were performed using TraceFinder software (Thermo Scientific).

For each class of lipids, the quantities of lipid species are given relative to their associated internal standard. Lipid notation follows that presented in Liebisch et al. (2020).

### Statistics

The statistical analysis and graphical representations were performed using Prism10 (GraphPad, Boston, MA, USA). Shapiro-Wilk normality test indicated that the relative abundances followed normal distributions for almost all lipid species. The ROUT test was used to identify outliers; outliers were identified only for a subset of fatty acids and all originated from the same brain. For each lipid species, com parisons of the three genotypes were performed using one-way ANOVA. Given that in most cases multiple ANOVA were performed for each lipid class, we controlled for the false discovery rate using the adaptive method of Benjamini, Krieger and Yekutieli with the FDR value Q set at 5%. When ANOVAs revealed significant influence of genotypes on lipid amounts (*q* < 5%), comparisons between pairs of genotypes were made using Holm-Šídák test, which allows correcting for multiple compar isons. To quantify the magnitude of differences between groups (Table 1), we calculated the ratio of the mean lipid species levels (expressed as percentage) of the 3 compared pairs: Akp2^−/−^/Akp2^+/+^, Akp2^−/−^/Akp2^+/–^ and Akp2^+/–^/Akp2^+/+^; the 95% confidence intervals of the ratios were calculated using Fieller’s exact formula (Motulsky 1995).

**Table 1.** List of phosphatidylcholine species whose abundance varied significantly across *Akp2* genotypes as revealed by one-way ANOVAs. "Species abundance ratio" (second column) is the amount of a given PC species expressed as a percentage of the sum of the amounts of PC species in *Akp2*^+/+^ mouse brains. Species were then attributed an "abundance rank" (third column) on the basis of their species abundance ratio. One-way ANOVAs revealed significant effect of genotype for 11/21 PC species (fourth column). These 11 PCs passed the FDR correction (*q* < 0.05, fifth column; “discoveries” correspond to ANOVA *P* values < 0.0437). Significant differences between pairs of genotypes (sixth column) are indicated in bold (Holm-Šídák posthoc test, seventh columns). The rightmost column presents the mean abundance ratios as a function of genotypes expressed as a percentage, with the 95% confidence interval given between brackets.

| PC Species | Species abundance ratio (%) | Abundance rank | *P* value one-way ANOVA | FDR *q* value | Compared genotypes | *P* value Holm-Šídák | Ratio (%) [95% confidence interval] |
|---|---|---|---|---|---|---|---|
| PC 32:0 | 20.0 | 2/21 | 0.004 | 0.0181 | **-/- vs. +/+** | **0.004** | **84.9 [77.3 – 93.4]** |
|  |  |  |  |  | **-/- vs. +/-** | **0.04** | **89.8 [83.0 – 97.0]** |
|  |  |  |  |  | +/- vs. +/+ | 0.2 | 94.5 [86.3 – 103.8] |
| PC 32:1 | 9.0 | 5/21 | 0.04 | 0.0468 | **-/- vs. +/+** | **0.04** | **87.2 [78.4 – 97.8]** |
|  |  |  |  |  | -/- vs. +/- | 0.2 | 92.3 [85.4 – 99.9] |
|  |  |  |  |  | +/- vs. +/+ | 0.2 | 94.5 [84.0 – 107.0] |
| PC 34:0 | 4.0 | 9/21 | 0.01 | 0.0214 | **-/- vs. +/+** | **0.01** | **87.7 [79.9 – 96.9]** |
|  |  |  |  |  | -/- vs. +/- | 0.1 | 94.2 [88.9 – 99.7] |
|  |  |  |  |  | +/- vs. +/+ | 0.1 | 93.2 [85.2 – 102.5] |
| PC 34:1 | 21.4 | 1/21 | 0.006 | 0.0189 | **-/- vs. +/+** | **0.005** | **86.9 [79.5 – 95.6]** |
|  |  |  |  |  | -/- vs. +/- | 0.1 | 93.3 [89.5 – 97.3] |
|  |  |  |  |  | +/- vs. +/+ | 0.1 | 93.1 [85.7 – 101.9] |
| PC 36:1 | 3.4 | 11/21 | 0.004 | 0.0181 | **-/- vs. +/+** | **0.003** | **88.0 [81.8 – 95.2]** |
|  |  |  |  |  | -/- vs. +/- | 0.1 | 93.0 [90.1 – 96.0] |
|  |  |  |  |  | +/- vs. +/+ | 0.1 | 94.7 [87.9 – 102.5] |
| PC 36:3 | 2.4 | 14/21 | 0.008 | 0.0191 | **-/- vs. +/+** | **0.007** | **87.4 [80.2 – 95.9]** |
|  |  |  |  |  | -/- vs. +/- | 0.1 | 92.9 [88.6 – 97.3] |
|  |  |  |  |  | +/- vs. +/+ | 0.1 | 94.2 [86,6 – 103,0] |
| PC 36:4 | 9.2 | 4/21 | 0.004 | 0.0181 | **-/- vs. +/+** | **0.004** | **83.9 [75.3 – 94.1]** |
|  |  |  |  |  | **-/- vs. +/-** | **0.04** | **89.2 [84.4 – 94.0]** |
|  |  |  |  |  | +/- vs. +/+ | 0.1 | 94.1 [85.4 – 104.5] |
| PC 38:3 | 1.6 | 15/21 | 0.03 | 0.034 | **-/- vs. +/+** | **0.03** | **88.3 [80.1 – 97.9]** |
|  |  |  |  |  | -/- vs. +/- | 0.2 | 94.2 [88.6 – 100.2] |
|  |  |  |  |  | +/- vs. +/+ | 0.2 | 93.7 [85.2 – 103.7] |
| PC 38:4 | 8.7 | 6/21 | 0.009 | 0.0191 | **-/- vs. +/+** | **0.008** | **85.4 [76.5 – 95.7]** |
|  |  |  |  |  | -/- vs. +/- | 0.1 | 91.3 [85.4 – 97.3] |
|  |  |  |  |  | +/- vs. +/+ | 0.1 | 93.5 [85.2 – 103.5] |
| PC 38:5 | 4.7 | 8/21 | 0.02 | 0.0214 | **-/- vs. +/+** | **0.01** | **84.8 [74.9 – 96.8]** |
|  |  |  |  |  | -/- vs. +/- | 0.2 | 91.9 [85.8 – 98.0] |
|  |  |  |  |  | +/- vs. +/+ | 0.2 | 92.3 [83.1 – 103.8] |
| PC 38:6 | 5.1 | 7/21 | 0.01 | 0.0214 | **-/- vs. +/+** | **0.02** | **88.3 [80.1 – 97.7]** |
|  |  |  |  |  | **-/- vs. +/-** | **0.04** | **90.2 [84.0 – 96.4]** |
|  |  |  |  |  | +/- vs. +/+ | 0.6 | 98.0 [90.3 – 106.8] |

## Results

In order to disclose consequences of TNAP dysfunction on brain lipid metabolism, we performed a lipidomic analysis on the brains of fifteen 7-day-old mice (5 *Akp2*^+/+^, 5 *Akp2*^+/-^ and 5 *Akp2*^-/-^ mice). Four major categories of lipids – fatty acids, neutral lipids, sphingolipids and glycerophospholipids – were examined.

Twenty-one straight-chain fatty acids species have been identified among 45 targeted (Figure 1). In con trol mice (*Akp2*^+/+^ mice), the most abundant fatty acid (FA) was palmitic acid (FA 16:0, 21% of all fatty acids). Stearic acid (FA 18:0) showed a nearly equal abundance (20%). The poly-unsaturated ω3 docosa hexaenoic acid (FA 22:6(n-3), DHA) and the poly-unsaturated ω6 arachidonic acid (FA 20:4(n-6)) both ranked at the third level in terms of abundance (14.8% each). Oleic acid (FA 18:1(n-9)) was the fifth most abundant fatty acid (12.7%). This ranking is fairly consistent with that reported in previous rodent studies (e.g., Baumann et al. 1968; Green et al. 1999; Demeilliers et al. 2017; Bai et al. 2020). The relative abundances of the different species of fatty acids did not differ between genotypes (one-way ANOVA, *P* > 0.05 for all comparisons, Figure 1).

**Figure 1:**
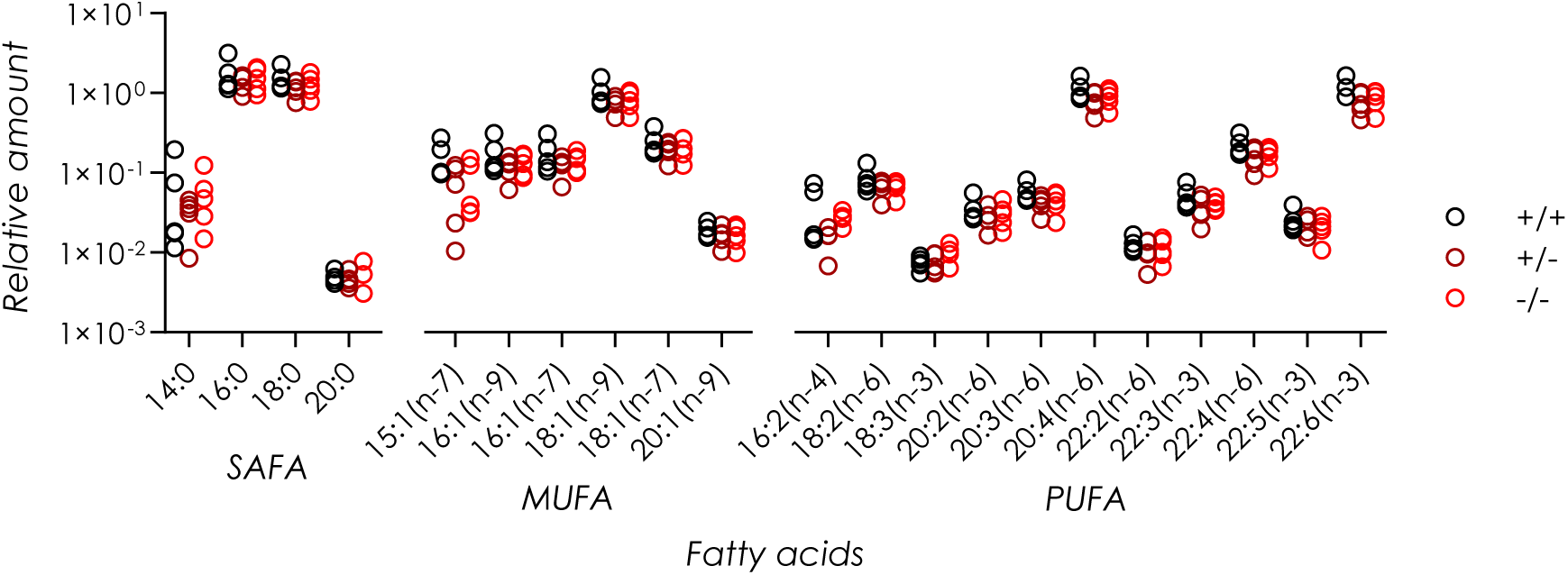
TNAP inactivation did not affect the total fatty acid (FA) content in the brains of 7-day-old mice. The amounts of each FA species are expressed relative to the internal standard TAG 19:0. Out of 15 targeted satu rated fatty acids (SAFA), 5 have been detected. Six monounsaturated fatty acids (MUFA) have been retrieved out of 11 targeted. Eleven polyunsaturated fatty acids (PUFA) have been detected out of 19 targeted. Outliers were identified for a subset of values, all originating from the same brain; the statistical outcome – no significant differences between genotypes – remained when these outliers were removed prior to analysis. Mouse genotype identified by dot color: black, *Akp2*^+/+^; dark red, *Akp2*^+/-^; red: *Akp2*^-/-^.

Neutral lipids classes examined in the present study included TAG and cholesterol. Six TAG species have been searched for but none were detected. TAG amounts were thus below our detection limits. This is consistent with previous studies that reported very low level of TAGs in rodent brain (Wells and Dittmer 1967; Chan et al. 2012; Jain et al. 2014). Diacylglycerols, also present at low level in rodent brain (Wells and Dittmer 1967; Chan et al. 2012; Jain et al. 2014; Fitzner et al. 2020), have not been looked for.

Cholesterol and three species of cholesteryl esters (CE 16, CE 18 and CE 20) have been looked for. Cholesterol and one species of cholesteryl esters (CE 16) have been detected in the three groups of mice. This is in agreement with previous studies that showed that, while cholesterol is very abundant in the brain, cholesteryl esters are generally present in very small quantities (Wells and Ditmer 1967; Chan et al. 2012; Jain et al. 2014; Fitzner et al. 2020). The relative abundances of cholesterol and CE 16 did not differ between genotypes (one-way ANOVA, *P* > 0.05 for both comparisons, Figures 2A and 2B).

**Figure 2:**
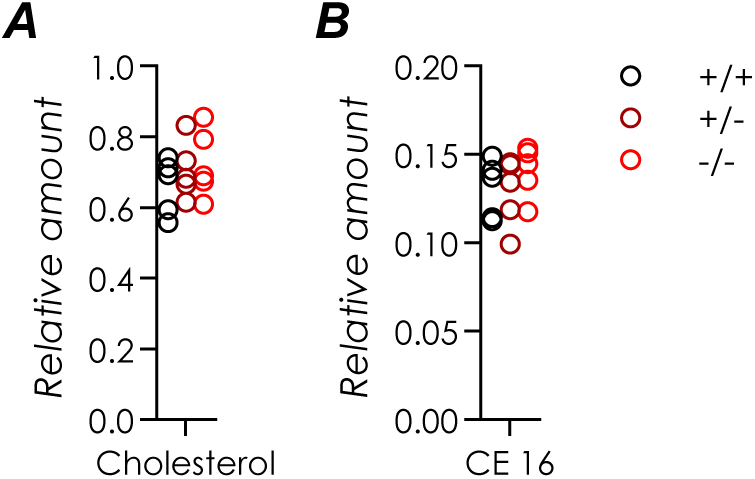
TNAP inactivation had no significant consequence on the abundance of neutral lipids in 7-day-old mouse brain. Among neutral lipids, triacylglycerols have not been detected. **A.** Abundance of cholesterol is expressed relative to the internal standard stigmasterol. **B.** The amount of C16cholesteryl ester (CE 16) is expressed relative to the internal standard CE 17:0. CE 16 was the sole cholesteryl ester species detected out of 3 targeted. Mouse genotype identified by dot color: black, *Akp2*^+/+^; dark red, *Akp2*^+/-^; red: *Akp2*^-/-^.

Eight sphingolipid classes have been scrutinized. Among these, 3 classes were not detected in our samples: dihydrosphingosine (sphinganine), sphingosine and gangliosides (GM1, GM2 and GM3 subclasses, 26 species targeted, none found). Sphinganine and sphingosine are intermediate in the metabo lism of sphingolipids and are present in small amounts relative to the other sphingolipids ( Davis et al. 2020), while the targeted gangliosides subclasses are only present at trace level in the central nervous system (Hilbig et al. 1982). The other classes (Figure 3) correspond to dihydroceramides (13 species targeted, one found, Figure 3A), ceramides (38 species targeted, 10 found, Figure 3B), monohexose ce ramides (36 species targeted, 5 found, Figure 3C), dihexose ceramides (9 species targeted, one found, Figure 3D) and sphingomyelin (41 species targeted, 18 found, Figure 3E). Within these various classes of sphingolipids, the most abundant species were systematically the 18:1;O2/18:0 (Figure 3A-D; labeled 36:1;O2 for sphingomyelins, Figure 3E). It alone represented 56% of ceramide relative abundance in *Akp2^+/+^*mice (Figure 3B) and 51% of that of sphingomyelins (Figure 3E), in agreement with previous studies (Chan et al. 2012; Jain et al. 2014; Fitzner et al. 2020). The relative abundances of the 35 sphingolipid species identified here were unaffected by TNAP inactivation (one-way ANOVA, *P* > 0.05 for all comparisons).

**Figure 3:**
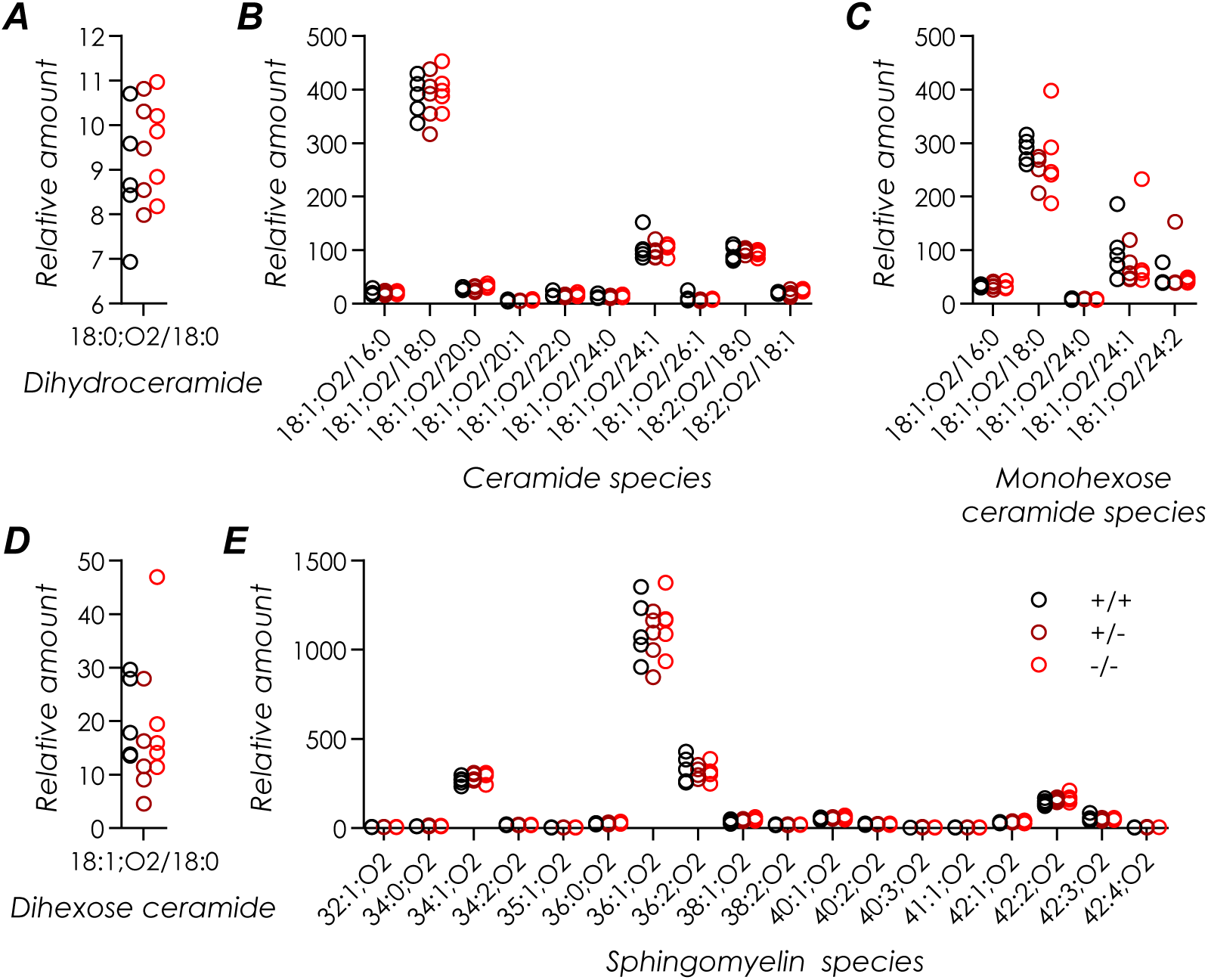
TNAP inactivation had no significant effect on mouse brain amounts of sphingolipids. **A.** Among the 13 species of dihydroceramide targeted, only dihydroceramide 18:0;O2/18:0 has been detected. Its relative abundance is expressed relative to the internal standard dihydroceramide 18:0;O2/12:0. **B.** Thirty eight species of ceramides have been targeted, of which 10 have been retrieved. Their relative abundances are expressed relative to the internal standard ceramide 18:1;O2/12:0. **C.** Five species of monohexose ceramides have been found out of 36 targeted. Their relative abundances are expressed relative to the internal standard glucosylceramide 18:1;O2/12:0. **D.** Out of 13 dihexose ceramide species targeted, only one has been detected. Its abundance is given relative to that of the internal standard lactosylceramide 18:1;O2/12:0. **E.** Eighteen different species of sphingomyelins have been detected among 41 targeted. Their relative abundances are given using sphingomyelin d18:1/12:0 as the internal standard. Mouse genotype identified by dot color: black, *Akp2*^+/+^; dark red, *Akp2*^+/-^; red: *Akp2*^-/-^.

Four classes of glycerophospholipids have been examined (Figure 4): phosphatidylserines (PS), phosphatidylinositols (PI), phosphatidylethanolamines (PE) and phosphatidylcholines (PC). Seventeen PS species have been detected, of which PS 40:6 was the most abundant (47.5% in *Akp2^+/+^* mice) followed by PS 40:5 (16.5%) (Figure 4A). Sixteen PI species were detected, in very low amounts except for PI 38:4, PI 38:3 and PI 38:5 which represented 57.5, 18 and 17.5% of the PIs (Figure 4B). Eighteen PE species have been detected, of which the most abundant were PE 38:4 and PE 38:3 (16% and 13%, re spectively) (Figure 4C). The PC class was the one with the greatest number of species (Figure 4D): 21 species, the most abundant being PC 34:1 (21.5%) and PC 32:0 (20%). Generally speaking, the relative abundances of the different glycerophospholipid species conform to that reported in previous rodent studies (e.g., Chan et al. 2012; Jain et al. 2014; Bascoul-Colombo et al. 2016; Fitzner et al. 2020).

**Figure 4:**
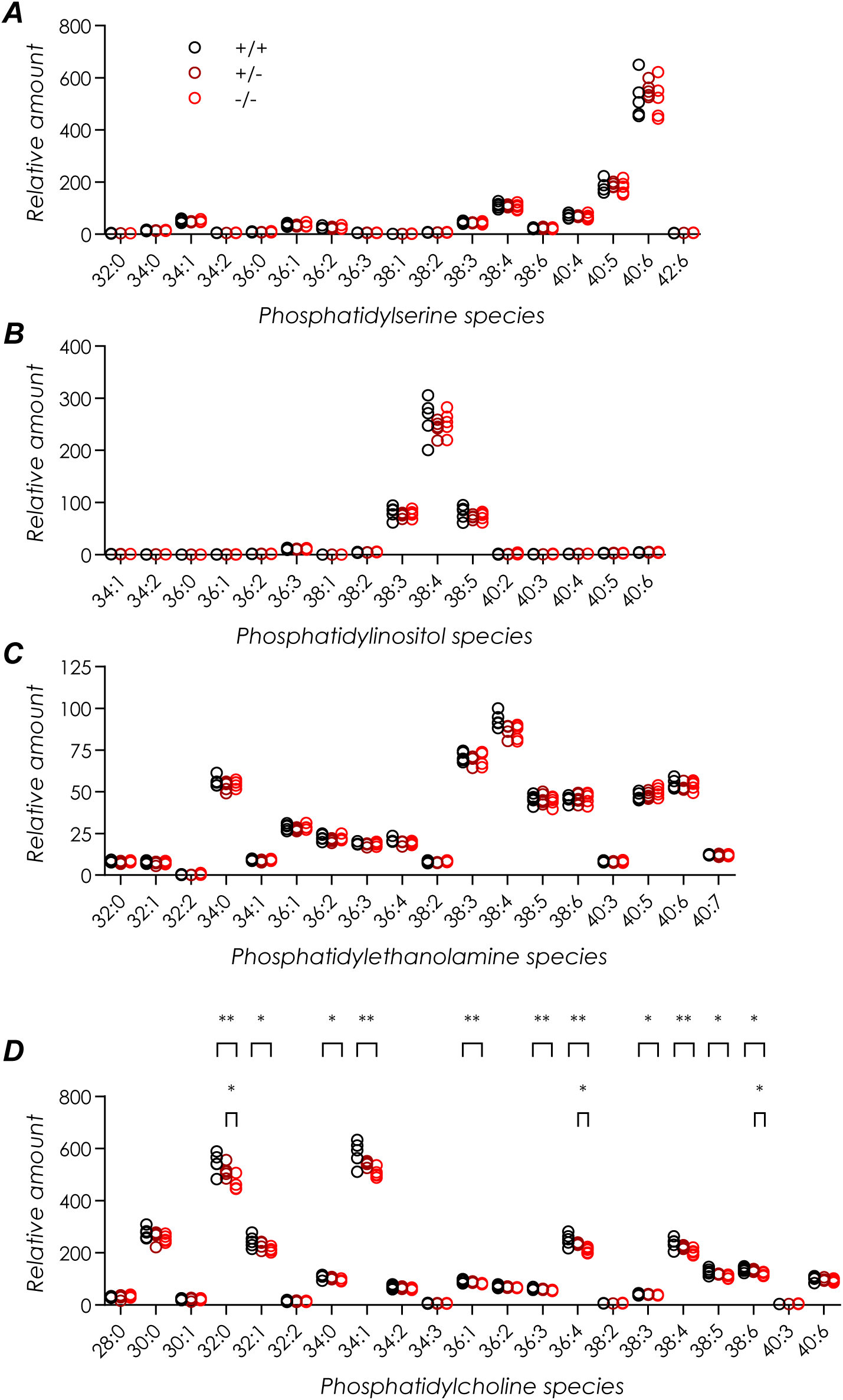
Among glycerophospholipids, TNAP inactivation significantly affected species belonging to the phosphatidylcholine class in the brains of 7-day-old mice. **A.** Eighteen species of phosphatidylserines (PSs) have been targeted, of which 17 have been retrieved. Their relative abundances is expressed relative to the internal standard PS 12:0/12:0. Presence or absence of functional TNAP did not affect the relative abundances of PSs. **B.** Out of 17 phosphatidylinositol (PI) species targeted, 16 have been detected. Their abundances are given relative to that of the internal standard PI 15:0/18:1. Their relative abundances was immune to the lack of functional TNAP. **C.** The 18 targeted species of phosphatidylethanolamine species (PE) have been detected. Their relative abundances are expressed relative to the internal standard PE 12:0/12:0. Genotype had no significant effect on PE species abundance. **D.** The 21 targeted species of phosphatidylcholine (PC) have been detected. The internal standard PC 13:0/13:0 has been used to compute their relative abundances. The abundance of 11 of the 21 PC species was significantly affected by mouse genotype (one-way ANOVA, threshold *P* value = 0.0437 after correcting for false discovery rate). The number of asterisks in **D** indicates significance level of paired comparisons using Holm-Šídák multiple comparison test: *: *P* < 0.05; **: *P* < 0.01. Mouse genotype identified by dot color: black, *Akp2*^+/+^; dark red, *Akp2*^+/-^; red: *Akp2*^-/-^.

The relative abundances of the PS and PI species showed no variation with respect to mouse genotypes (one-way ANOVA, *P* > 0.05 for all comparisons; Figures 4A and 4B). Mouse genotype seemed to have a significant influence on the amount of two species of the PE class (one-way ANOVA, *P* = 0.0411 and 0.0433; Figure 4C). However these differences were dismissed after controlling for the false discovery rate (*q* = 0.34 in both cases).

On the other hand, the amounts of species of the PC class appeared to be significantly influenced by the genotype (one was ANOVA, *q* < 0.05, FDR-adjusted *P* value < 0.0437). These results are summarized in Table 1 and in Figure 4D. Significant effect of genotype was observed for 11 species, including 8 of the 9 most abundant species (Figure 4D, Table 1). Pairwise comparisons revealed significant differences between *Akp2*^-/-^ and *Akp2*^+/+^ genotypes in these 11 cases (Table 1, Figure 4D). Significant differences were also observed between *Akp2*^-/-^ and *Akp2*^+/-^ genotypes in 3 of these 11 cases (Table 1, Figure 4D). The 11 species that showed significant difference between *Akp2*^-/-^ and *Akp2*^+/+^ mice represented 89.5% of the total PC abundance in *Akp2*^+/+^ mice. When compared to *Akp2*^+/+^ mice, species amounts were reduced by 11.7% to 16.1% in *Akp2*^-/-^ mice (Table 1) and by 13.9% on average (weighted average for all species with significant decreases). Differences between *Akp2*^+/-^ and *Akp2*^+/+^ were intermediate between those observed between *Akp2*^-/-^ and *Akp2*^+/+^ mice (Table 1).

## Discussion

### Inactivation of TNAP does not significantly alter brain levels of sphingolipids in 7-day-old mice

An increase in serum PLP level is one of the hallmarks of hypophosphatasia in human (Whyte et al. 1985) as well as in mouse models of hypophosphatasia (Waymire et al. 1995; Fedde et al. 1999). The PLP level was also found to be considerably elevated (5 to 10 times more than the normal range) in the CSF of a child with severe hypophosphatasia before pyridoxine treatment (van der Stoep et al. 2014). In TNAP KO mice, compromised dephosphorylation of extracellular PLP results in a *reduction* in intracellular PLP concentration (Waymire et al. 1995). PLP is a cofactor of serine palmitoyltransferase, the enzyme responsible for the first step of sphingolipid synthesis (Braun et al. 1970; Hanada 2003; Bourquin et al. 2011). In accordance with this PLP dependency, studies on consequences of vitamin B6 deprivation reported decreased amounts of sphingolipids in the brains of young adult rodents (Williamson and Coniglio 1971; Kurtz et al. 1972; Kurtz and Kanfer 1973). These observations predicted that there would also be a decrease in the amount of all sphingolipid species in the brains of TNAP KO mice. Yet our results failed to support this prediction.

Animal age may offer a first explanation for this negative results. Indeed, Williamson and Coniglio (1971) reported that the effect of vitamin B6 deprivation in rats depended on the age of the animals: although the amounts of various brain sphingolipids were decreased at postnatal day (PND) 21, vitamin B6 deprivation had no effect on sphingomyelins, cerebrosides, ceramides, and sulfatides at PND 14 and PND 7. Our PND 7 mice would be in the latter case.

It is also worth noting that PLP and PL are not completely absent from the brain of TNAP KO mice. In mice aged 10-14 days, PLP and PL amounts in the brains of *Akp2*^-/-^ mice are at 35% and 60%, respectively, of the levels in *Akp2*^+/+^ mice (Waymire et al. 1995). Part of this remnant PL/PLP may be derived from mother milk, which is strongly enriched in vitamin B6 (Chang and Kirksey 2002; Oosterink et al. 2015). Maintenance of sphingolipid synthesis despite reduced brain level of PLP at PND 7 suggests that the affinity of serine palmitoyltransferase for PLP is relatively high (Cf. Braun et al. 1970).

### Myelination deficits in TNAP KO mice do not translate into detectable myelin lipid changes: NAA hypothesis refuted?

Fonta et al. (2005) showed that, during development, axons undergoing myelination display a particularly high alkaline phosphatase activity. This raised the possibility that TNAP is involved in myelination. This possibility was supported by the observation of defective myelination in the spinal cord and cerebral cortex of 7to 10-day-old TNAP KO mice (Hanics et al. 2012). This defective myelination may be due to vitamin B6 defective metabolism; indeed, vitamin B6 deprivation leads to a decrease in the number of myelinated axons of the pyramidal tract in 15-day-old rats (Morré et al. 1978) and to a reduction in myelin amount in 20-day-old rats (Kurtz and Kanfer 1973). Alternatively, this may be a consequence of abnormalities in nucleotide metabolism: Kermer et al. (2010) showed that TNAP inhibition impairs the differentiation of precursor cells into oligodendrocytes; oligodendrocyte differentiation and matu ration are promoted by adenosine (Stevens et al. 2002); and adenosine levels are markedly reduced in the brains of TNAP-KO mice (Cruz et al. 2017). Regardless of the mechanisms, our previous study also revealed increased amounts of NAA in the brains of TNAP KO mice (Cruz et al. 2017). As NAA has been suggested to be involved in myelin lipid production (Namboodiri et al. 2006), this led us to hypothesize that the increased amount of NAA observed in the brains of Akp2^-/-^ mice resulted from reduced production of myelin lipids.

The decrease in PCs we observed is unlikely to be a consequence of defective myelination. This is because the process of myelination is associated with a decrease in the proportion of PC in myelin (Horrocks 1968; Norton and Poduslo 1973; Naffaa et al. 2022). Thus, defective myelination should result in an increase of the relative proportion of PC in the brains of *Akp2^-/-^* mice.

Besides, the present study failed to reveal significant changes in myelin-associated lipids. In adult, myelin is particularly enriched in monohexose ceramides (cerebrosides) (O’Brien and Sampson 1965a; Kishimoto et al. 1969; Ghandour and Skoff 1988; Camargo et al. 2017; Fitzner et al. 2020; Naffaa et al. 2022; Osetrova et al. 2024). Yet amounts of monohexose ceramides did not differ between *Akp2*^-/-^ and *Akp2*^+/+^ mice^1^.

Two basic facts help mitigate our negative result. First, although some myelinated fibers can be observed in the cortex of 7to 10-day-old *Akp2*^+/+^ mice (Hanics et al. 2012), myelin formation in the rodent brain^2^ is barely initiated at this age (Schonbach et al. 1968; Norton and Poduslo 1973; Shiota et al. 1989; Davis et al. 2020; Naffaa et al. 2022). Based on available data (Norton and Poduslo 1973), we can roughly estimate that myelin lipids would account for less than a couple percents of the total lipids in the brains of 7-day-old mice^3^. Second, myelin-specific lipids appear quite late in development (Wells and Dittmer 1967; Horrocks 1968; Davis et al. 2020). In particular, hexose ceramides are barely detectable in oligodendrocyte at PND 8 (Davis et al. 2020) while in myelin, the amounts of hexose ceramides increase sharply only after PND 15 (Horrocks 1968; Norton and Poduslo 1973; Naffaa et al. 2022). It is also noticeable that, in the present study, the myelin-specific fatty acids FA 20:1, FA 24:0 and FA 24:1 (O’Brien and Sampson 1965b; Kishimoto et al. 1969; Schmitt et al. 2015) were either not detected (FA 24:0, FA 24:1) or present in trace amount (FA 20:1, 0.3% of the detected fatty acids, to be compared with 2.4% in 7to 10-week old mice in Baumann et al. 1968). Therefore, even if defective myelination had led to changes in the amounts of myelin-specific lipid species in *Akp2*^-/-^ mice, these changes were probably too small in magnitude to be detectable in our analysis of whole brain lipids.

The >10% increases in NAA we previously reported in TNAP KO mice is therefore difficult to recon cile with a barely begun myelination and lack of detectable change in the amount of myelin-specific lipids in these mice. While NAA is synthesized mainly by neurons, its catabolism is exclusively performed by oligodendrocytes. The enzyme responsible for this catabolism is aspartoacylase, an enzyme that is only expressed in oligodendrocytes (Namboodiri et al. 2006; Grønbæk-Thygesen and HartmannPetersen 2024). Aspartoacylase activity is strong in mature oligodendrocytes, but is also present, albeit at a lower level, in non myelinating, immature oligodendrocytes (Bhakoo et al. 2001). As inhibition of TNAP results in a drastic impairment of the differentiation of progenitor cells into oligodendrocytes (Kermer et al. 2010), aspartoacylase activity was likely to be lower in TNAP KO mice, thereby leading to increased concentration of NAA. Yet NAA accumulation and defective myelination would have no causal relationship and would be independent consequences of deficits in oligodendrocytes differentiation and maturation.

### Among glycerophospholipids, only phosphatidylcholines are affected

The main result of our study is that TNAP dysfunction results in a reduction in the brain amounts of PCs. This reduction was observed in approximately half of the PC species and ranged from 11.7% to 16.1%. The affected PC species were those that were the most abundant, representing 89.5% of the to tal amount of PCs.

A first explanation for the reduced amount of PCs would simply be that there is a reduction in the rate of growth of brain cells in *Akp2*^-/-^ mice. Yet, if the main difference between *Akp2*^-/-^ and wild type mice was simply a decrease in the requirements for lipid production in *Akp2*^-/-^ mice, then all lipids would be affected equally, as neuron differentiation and brain development are accompanied by an increase in concentration for most glycerophospholipid and sphingolipid species (Wells and Dittmer 1967; Williamson and Coniglio 1971; Araki and Wurtman 1997; Marcucci et al. 2010; Camdzic et al. 2023). Our results, instead, showed no significant variation for lipid species outside of the PC class; in 7-dayold mice, TNAP dysfunction seems to specifically affect PCs.

PCs can be synthesized through two pathways (reviewed in: Farooqui et al. 2000a; van der Veen et al. 2017; Tavasoli et al. 2020). The first pathway is called the Kennedy pathway (Kennedy and Weiss 1956) and is the main pathway for PC synthesis. It uses diacylglycerols and choline as precursors. The second pathway for PC synthesis, Bremer pathway, (Bremer et al. 1960; reviewed in: Farooqui et al. 2000a; van der Veen et al. 2017) uses PEs as precursor, which are methylated three times using S-adenosyl methionine (SAM) as methyl donor. This triple methylation is catalyzed by PEMT, which is encoded by *Pemt* gene (Walkey et al. 1997).

One explanation for the specific decrease in PC levels could be a reduction in the amount of choline in the brains of Akp2^-/-^ mice, which would impede PC production through the Kennedy pathway. Choline is an essential nutrient that is largely provided by the diet (Blusztajn et al. 2017; Wallace et al. 2020), although *de novo* synthesis through the PE methylation pathway can partially compensate for choline deficiency (van der Veen et al. 2017; Wallace et al. 2020). It could be envisioned that inactivation of TNAP altered choline absorption, metabolism or transport downstream the brain of *Akp2*^-/-^ mice. Yet, as far as the brain is concerned, Cruz et al. (2017) showed that choline level in *Akp2*^-/-^ mice does not differ from that in *Akp2*^+/+^; this implies that choline deficiency was not responsible for the decrease in the amount of PCs in the brains of TNAP KO mice.

A second possible explanation for the decrease in PC levels we observed lies on the indirect depen dency of the second PC synthesis pathway, Bremer pathway, on PLP availability. Kurtz et al. (1972) showed that vitamin B6 deprivation results in a decrease (-15%) in the amount of brain glycerophospholipids in 18-day-old rats. More detailed studies (Suzuki and Okada 1982; Loo and Smith 1986; She et al. 1995), which examined the consequences of vitamin B6 deprivation in the liver, reported a specific decrease in the amount of PCs in this condition. Two of these studies (Loo and Smith 1986; She et al. 1995) further showed that vitamin B6 deprivation resulted in a parallel increase in S-adenosyl homocysteine (SAH) concentration (see also Nguyen et al. 2001; Isa et al. 2006a). This increase in SAH concentration is the consequence of the strong dependency of the methionine cycle and transsulfuration pathway on PLP availability (Gregory et al. 2016). Due to a shortage of intracellular PLP in TNAP KO mouse brain (Waymire et al. 1995), the transsulfuration pathway and methionine cycle are also dysfunctional, as evidenced by the accumulation of cystathionine and methionine (Cruz et al. 2017^4^; Phoenix et al. 2025). Reduced PLP availability results in an impairment of the transsulfuration pathway, leading to increased homocysteine concentration (Martinez et al. 2000; Isa et al. 2006a; Kitagawa et al. 2015). Under physiological conditions, homocysteine, along with adenosine, is the breakdown product of SAH. However, in the presence of increased levels of homocysteine, the reaction is reversed such that SAH is then produced from homocysteine and adenosine. (Reddington and Pusch 1983; Choumenkovitch et al. 2002; Kloor and Osswald 2004; Isa et al. 2006b; Kitagawa et al. 2015). The increase in the amount of SAH can compromise PEMT activity. Indeed, PEMT, similarly to many other methyl transferases, is inhibited by SAH (Blusztajn et al. 1979; Hoffman et al. 1981; She et al. 1995). In short, we suggest that the decrease in PC level that we observed in TNAP KO mice results from an in hibition of PEMT due to an increased level in SAH, the latter being the consequence of a dysfunction of the methionine cycle, itself the consequence of a blockage of the transsulfuration pathway due to decrease in intracellular PLP level.

Presence of PEMT activity in the brain has been reported in a considerable number of studies (Chida and Arakawa 1971; Blusztajn et al. 1979; Tashiro et al. 1983; Cornatzer et al. 1984; Blusztajn et al. 1985; Lakher and Wurtman 1987; Hargreaves and Clandinin 1989; Johnson and Blusztajn 1998; Guan et al. 1999; Marcucci et al. 2010), even though it has been reported to be negligible in some specific neuron subtypes (NG108-15 neuroblastoma: de Blas et al. 1984; sympathetic neurons and PC12 cells: Vance et al. 1994), suggesting that PEMT expression may be cell type specific. The mRNA encoding for PEMT and the corresponding protein have been localized in the brain of both mice, pigs and humans ( Sjöstedt et al. 2020: the HPA Brain Atlas, www.proteinatlas.org/brain). In the adult, PEMT activity and mRNA expression level appear to be lower in the brain compared to the liver (Chida and Arakawa 1971; Blusztajn et al. 1979; Cornatzer et al. 1984; Sjöstedt et al. 2020). However this is also the case for choline phosphotransferase (PCT) activity of the Kennedy pathway, such that the ratio of PEMT/PCT activities may actually be higher in the brain than in the liver (Cornatzer et al. 1984). In addition, activity and expression of PEMT are developmentally regulated and appear to be quite higher in the brain of mice and rats aged a few days compared to adults (Tashiro et al. 1983; Blusztajn et al. 1985; Naffaa et al. 2022). Of further interest, Zhu et al. 2004 found that PC amounts in the brains of *Pemt* KO mice was reduced by 19% in comparison to that in the corresponding wildtype mice, a decrease quite close to that reported in the present study. A possible argument against our PEMT-inhibition-by-SAH hypothesis is that PEMT may be preferentially involved in the production of long-chain polyunsaturated PC species (DeLong et al. 1999; Pynn et al. 2011). Yet our results showed significant reductions in PC amounts independently of their acyl chain length and saturation level. It should be noticed, however, that phospholipids undergo extensive remodeling by deacylation and reacylation, a process known as the Lands cycle (reviewed in Farooqui et al. 2000b; O’Donnell 2022; Valentine et al. 2023), which results in continuous shuttling of fatty acids between different phospholipid species. Thus, it has been shown that PCs synthesized by PEMT can be remodeled such that, in the long term (≥ 24 h), their chain lengths and saturation levels equilibrate with those of PCs synthesized by the Kennedy pathway (Pynn et al. 2011).

### Consequences of altered phosphatidylcholine levels

Decrease in PC production has many detrimental consequences. This issue has been extensively studied with respect to liver physiology (reviewed in van der Veen et al. 2017; Wallace et al. 2020). In the liver, even small decreases in the amount of PC, or in the PC/PE ratio, lead to alterations in lipoprotein secretion leading to hepatic steatosis as well as disruption of the integrity of cell membranes.

During brain development, neuronal differentiation is associated with an increase in PC production (Marcucci et al. 2010). It has also been shown that increasing the amount of PC, whether through exogenous supply or stimulation of synthesis, can by itself promote neuronal differentiation ( Marcucci et al. 2010; Paoletti et al. 2016; Montaner et al. 2018). Furthermore, studies have shown that inhibiting PC synthesis in neuronal cell cultures reduces or suppresses axonal growth, and this occurs even if the production of other glycerophospholipids (PE and PS) is within the normal range (Posse de Chaves et al. 1995a, 1995b). As TNAP dysfunction specifically reduces PC level, we could expect that axon growth may also be compromised in *Akp2*^-/-^ mice. Although axon growth depends on multiple parameters, it is interesting to note that studies, also using the *Akp2* mouse model, have indeed reported a decrease in the density of axons in the white matter of the spinal cord (Hanics et al. 2012), as well as a reduction in the length of the axons of cortical neurons that have interhemispheric projections via the corpus callo sum (Sebastián-Serranoa et al. 2016).

Glycerophospholipid composition and metabolism appear to be abnormal in a number of neurodegen erative diseases (reviewed in Farooqui et al. 2000a; Wood 2012; Xicoy et al. 2019). In particular, Alzheimer’s disease is associated with a decrease in the amounts of choline and ethanolamine, the PC and PE precursors, and an increase in the amounts of glycerophosphocholine and glycerophospho ethanolamine, the degradation products of PC and PE. (Nitsch et al. 1992; Wood 2012; Batra et al. 2022). Decreased levels of PCs, PEs and PSs have been observed in specific brain regions of Hunting ton’s disease patients (Phillips et al. 2022). Parkinson’s disease also is characterized by altered levels of PE and PC (Xicoy et al. 2019). Besides, studies showed that the dysfunction of enzymes involved in the Kennedy pathway leads to neurodevelopmental disorders (Mitsuhashi et al. 2011; Hoover-Fong et al. 2014; Klöckner et al. 2022). More generally, altered or dysfunctional lipid metabolism is associated with several brain diseases and neurological disorders (reviewed in Lauwers et al. 2016; Harayama and Riez man 2018; Tavasoli et al. 2020; Rickman et al. 2020; Yoon et al. 2022). Our study thus adds hypophosphatasia to the list of diseases associated with changes in cerebral lipid metabolism.

## Acknowledgments

This study was supported by a “Tremplin” grant from University Paul Sabatier (Toulouse, 2022): Chimie analytique pour l’étude des Fonctions hépatiques de la phosphatase alcaLINE (FéLINE). José Luis Millán is warmly thanked for the gift of the *Akp2* mice that were used to establish our breeding colony. Thomas Cruz is thanked for his participation in the early stages of the metabolomic studies. Lipidomic analysis was performed at MetaToul (Toulouse metabolomics & fluxomics facilities, www.mth-metatoul.com, on the site dedicated to Lipids Analysis at Inserm I2MC) and was supported by the MetaboHUB infrastructure funded by the Agence Nationale de la Recherche under the France 2030 program (MetaboHUB ANR-11-INBS-0010; MetEx + ANR-21-ESRE-0035; MetaboHUB (JVCE) ANR-24-INBS-0012).

## Authors contribution

Conceptualization: LGN, SB, VG, JBM, CF

Investigation: MG, SB, OD, EM

Data Curation: LGN, OD.

Formal Analysis: LGN Visualization: LGN

Writing – Original Draft Preparation: LGN

Writing – Review & Editing: LGN, MG, SB, OD, VG, JBM, CF

Funding Acquisition: LGN, VG, JBM, CF

Project Administration: LGN, VG, JBM, CF

Supervision: LGN, VG, JBM, CF

## Abbreviations

AP: alkaline phosphatase
FA: fatty acid
FDR: false discovery rate
NAA: N-acetylaspartate
NAAG: N-acetyl-aspartylglutamate
PC: phosphatidylcholine
PE: phosphatidylethanolamine
PEA: phosphoethanolamine
PEMT: phosphatidylethanolamine methyl transferase
PI: phosphatidylinositol
PL: pyridoxal
PLP: pyridoxal phosphate
PND: postnatal day
PPi: pyrophosphate
PS: phosphatidylserine
SAH: S-adenosyl homocysteine
SAM: S-adenosyl methionine TAG: triacylglycerol
TNAP: tissue-nonspecific alkaline phosphatase

## Footnotes

1 The gangliosides that were targeted (and not detected) in the present study (GM1, 2 & 3) are not specific to white matter; the ones that are found on myelinated axons are GD1a and GT1b but these were not examined.

2 The spinal cord, where myelination is more advanced at PND 7 than in the brain, was not examined in the present study.

3 Norton and Poduslo (1973) estimated that myelin represents 0.3% of the dry weight of the rat brain at PND 10. At the same age, total lipids represent 27.7% of the dry weight. Assuming myelin is predominantly composed of lipids, then myelin lipids would represent about 1% of the total lipids. Supposing that their centrifugation method did not isolate all the myelin, we propose a value of 2% to include a safety margin.

4 The other metabolites of the transsulfuration pathway and methionine cycle – SAM, SAH, homocysteine and cysteine – were not detected due to low sensitivity of the untargeted ^1^H-NMR method used in Cruz et al.’s study.

## References

1. Araki W, Wurtman RJ (1997) Control of membrane phosphatidylcholine biosynthesis by diacylglycerol levels in neuronal cells undergoing neurite outgrowth. Proc Natl Acad Sci U S A 94: 11946–11950. doi: 10.1073/pnas.94.22.11946. PMID: 9342342; PMCID: PMC23665.

2. Bai YR, Wang YY, Meng DL, Shi ZL, Song XF, Yang ZZ, Zhang W, Yang L, Shi RL (2020) Alterations of fatty acid composition and metabolism in APP/PS1 transgenic mice. Neurosci Lett 738: 135401. doi: 10.1016/j.neulet.2020.135401. PMID: 32976920.

3. Barrans A, Collet X, Barbaras R, Jaspard B, Manent J, Vieu C, Chap H, Perret B (1994) Hepatic lipase induces the formation of pre-beta 1 high density lipoprotein (HDL) from triacylglycerol-rich HDL2. A study comparing liver perfusion to in vitro incubation with lipases. J Biol Chem 269: 11572–11577. PMID: 8157689.

4. Bascoul-Colombo C, Guschina IA, Maskrey BH, Good M, O’Donnell VB, Harwood JL (2016) Dietary DHA supplementation causes selective changes in phospholipids from different brain regions in both wild type mice and the Tg2576 mouse model of Alzheimer’s disease. Biochim Biophys Acta 1861: 524–537. doi: 10.1016/j.b-balip.2016.03.005. PMID: 26968097; PMCID: PMC4847476.

5. Baslow MH, Guilfoyle DN (2013) Canavan disease, a rare early-onset human spongiform leukodystrophy: insights into its genesis and possible clinical interventions. Biochimie 95: 946–956. doi: 10.1016/j.biochi.2012.10.023. PMID: 23151389.

6. Batra R, Arnold M, Wörheide MA, Allen M, Wang X, Blach C, Levey AI, Seyfried NT, Ertekin-Taner N, Bennett DA, Kastenmüller G, Kaddurah-Daouk RF, Krumsiek J; Alzheimer’s Disease Metabolomics Consortium (ADMC) (2022). The landscape of metabolic brain alterations in Alzheimer’s disease. Alzheimers Dement 19: 980–998. doi: 10.1002/alz.12714. PMID: 35829654; PMCID: PMC9837312.

7. Baumann NA, Jacque CM, Pollet SA, Harpin ML (1968) Fatty acid and lipid composition of the brain of a myelin deficient mutant, the "quaking" mouse. Eur J Biochem 4: 340–344. doi: 10.1111/j.1432-1033.1968.tb00216.x. PMID: 5653767.

8. Baumgartner-Sigl S, Haberlandt E, Mumm S, Scholl-Bürgi S, Sergi C, Ryan L, Ericson KL, Whyte MP, Högler W (2007) Pyridoxine-responsive seizures as the first symptom of infantile hypophosphatasia caused by two novel missense mutations (c.677T>C, p.M226T; c.1112C>T, p.T371I) of the tissue-nonspecific alkaline phosphatase gene. Bone 40: 1655–1661. doi: 10.1016/j.bone.2007.01.020.

9. Beckonert O, Keun HC, Ebbels TMD, Bundy J, Holmes E, Lindon JC, Nicholson JK (2007) Metabolic profiling, metabolomic and metabonomic procedures for NMR spectroscopy of urine, plasma, serum and tissue extracts. Nat Protoc 2: 2692–2703. doi: 10.1038/nprot.2007.376. PMID: 18007604.

10. Bhakoo KK, Craig TJ, Styles P (2001) Developmental and regional distribution of aspartoacylase in rat brain tissue. J Neurochem 79: 211–220. doi: 10.1046/j.1471-4159.2001.00561.x. PMID: 11595773.

11. Bourquin F, Capitani G, Grütter MG (2011) PLP-dependent enzymes as entry and exit gates of sphingolipid metabolism. Protein Sci 20: 1492–1508. doi: 10.1002/pro.679.

12. Bowden SA, Foster BL (2019) Alkaline phosphatase replacement therapy for hypophosphatasia in development and practice. In: advances in experimental medicine and biology volume 1148: therapeutic enzymes: function and clinical implications, Labrou N. (ed), Springer, Singapore, pp. 279–322. doi: 10.1007/978-981-13-7709-9_13.

13. Braun PE, Morell P, Radin NS (1970) Synthesis of C_18_and C_20_-dihydrosphingosines, ketodihydrosphingosines, and ceramides by microsomal preparations from mouse brain. J Biol Chem 245: 335–341. PMID: 4391620.

14. Bremer J, Figard PH, Greenberg DM (1960) The biosynthesis of choline and its relation to phospholipid metabolism. Biochim biophys acta 43: 477–488. doi: 10.1016/0006-3002(60)90470-4.

15. Blusztajn JK, Zeisel SH, Wurtman RJ (1979) Synthesis of lecithin (phosphatidylcholine) from phosphatidylethanolamine in bovine brain. Brain Res 179: 319–327. doi: 10.1016/0006-8993(79)90447-5. PMID: 509240.

16. Blusztajn JK, Zeisel SH, Wurtman RJ (1985) Developmental changes in the activity of phosphatidylethanolamine N-methyltransferases in rat brain. Biochem J 232: 505–511. doi: 10.1042/bj2320505. PMID: 4091805; PMCID: PMC1152909.

17. Blusztajn JK, Slack BE, Mellott TJ (2017) Neuroprotective actions of dietary choline. Nutrients 9: 815. doi: 10.3390/nu9080815. PMID: 28788094; PMCID: PMC5579609.

18. Camargo N, Goudriaan A, van Deijk AF, Otte WM, Brouwers JF, Lodder H, Gutmann DH, Nave KA, Dijkhuizen RM, Mansvelder HD, Chrast R, Smit AB, Verheijen MHG (2017) Oligodendroglial myelination requires astrocyte-derived lipids. PLoS Biol 15: e1002605. doi: 10.1371/journal.pbio.1002605. PMID: 28549068; PMCID: PMC5446120.

19. Camdzic M, Aga DS, Atilla-Gokcumen GE (2023) Cellular lipidome changes during retinoic acid (RA)-induced differentiation in SH-SY5Y cells: a comprehensive *in vitro* model for assessing neurotoxicity of contaminants. Environ Health (Wash) 1: 110–120. doi: 10.1021/envhealth.3c00022. PMID: 37614295; PMCID: PMC10443778.

20. Chan RB, Oliveira TG, Cortes EP, Honig LS, Duff KE, Small SA, Wenk MR, Shui G, Di Paolo G (2012) Comparative lipidomic analysis of mouse and human brain with Alzheimer disease. J Biol Chem 287: 2678–2688. doi: 10.1074/jbc.M111.274142. PMID: 22134919; PMCID: PMC3268426.

21. Chang SJ, Kirksey A (2002) Vitamin B6 status of breast-fed infants in relation to pyridoxine HCl supplementation of mothers. J Nutr Sci Vitaminol (Tokyo) 48: 10–17. doi: 10.3177/jnsv.48.10. PMID: 12026182.

22. Chida N, Arakawa T (1971) Metabolism of phosphatidylcholine in brain and liver of developing rats. Tohoku J Exp Med 104: 359–371. doi: 10.1620/tjem.104.359. PMID: 5124684.

23. Choumenkovitch SF, Selhub J, Bagley PJ, Maeda N, Nadeau MR, Smith DE, Choi SW (2002) In the cystathionine beta-synthase knockout mouse, elevations in total plasma homocysteine increase tissue S-adenosylhomocysteine, but responses of S-adenosylmethionine and DNA methylation are tissue specific. J Nutr 132: 2157–2160. doi: 10.1093/jn/132.8.2157.

24. Coburn SP (2015) Vitamin B-6 Metabolism and Interactions with TNAP. Subcell Biochem 76: 207–238. doi: 10.1007/978-94-017-7197-9_11. PMID: 26219714.

25. Colazo JM, Hu JR, Dahir KM, Simmons JH (2019) Neurological symptoms in Hypophosphatasia. Osteoporos Int 30: 469–480. doi: 10.1007/s00198-018-4714-3. PMID: 30215116.

26. Connors BW, Malenka RC, Silva LR (1988) Two inhibitory postsynaptic potentials, and GABA_A_ and GABA_B_ receptor-mediated responses in neocortex of rat and cat. J Physiol 406: 443–468. doi: 10.1113/jphysiol.1988.sp017390.

27. Cornatzer WE, Haning JA, Wallwork JC, Sandstead HH (1984) Effect of zinc deficiency on the biosynthesis of phosphatidylcholine in rat microsomes. Biol Trace Elem Res 6: 393–401. doi: 10.1007/BF02989256. PMID: 24264176.

28. Cruz T, Gleizes M, Balayssac S, Mornet E, Marsal G, Millán JL, Malet-Martino M, Nowak LG, Gilard V, Fonta C (2017) Identification of altered brain metabolites associated with TNAP activity in a mouse model of hypophosphatasia using untargeted NMR-based metabolomics analysis. J Neurochem 140: 919–940. doi: 10.1111/jnc.13950. PMID: 28072448; PMCID: PMC5339068.

29. Davis DL, Mahawar U, Pope VS, Allegood J, Sato-Bigbee C, Wattenberg BW (2020) Dynamics of sphingolipids and the serine palmitoyltransferase complex in rat oligodendrocytes during myelination. J Lipid Res 61: 505–522. doi: 10.1194/jlr.RA120000627. PMID: 32041816; PMCID: PMC7112141.

30. de Blas A, Adler M, Shih M, Chiang PK, Cantoni GL, Nirenberg M (1984) Inhibitors of CDP-choline synthesis, action potential calcium channels, and stimulus-secretion coupling. Proc Natl Acad Sci USA 81: 4353–4357. doi: 10.1073/pnas.81.14.4353. PMID: 6087319; PMCID: PMC345587.

31. de Roo MG, Abeling NG, Majoie CB, Bosch AM, Koelman JH, Cobben JM, Duran M, Poll-The BT (2014) Infantile hypophosphatasia without bone deformities presenting with severe pyridoxine-resistant seizures. Mol Genet Metab 111: 404–407. doi: 10.1016/j.ymgme.2013.09.014. PMID: 24100244.

32. DeLong CJ, Shen YJ, Thomas MJ, Cui Z (1999) Molecular distinction of phosphatidylcholine synthesis between the CDP-choline pathway and phosphatidylethanolamine methylation pathway. J Biol Chem 274: 29683–29688. doi: 10.1074/jbc.274.42.29683. PMID: 10514439.

33. Demeilliers C, Toufektsian MC, Salen P, Laporte F, Petroni K, de Lorgeril M (2017) Ethanol drinking, brain mitochondrial DNA, polyunsaturated fatty acids and effects of dietary anthocyanins. Clin Nutr Exp 12: 11–19.

34. Demirbilek H, Alanay Y, Alikaşifoğlu A, Topçu M, Mornet E, Gönç N, Özön A, Kandemir N (2012) Hypophosphatasia presenting with pyridoxine-responsive seizures, hypercalcemia, and pseudotumor cerebri: case report. J Clin Res Pediatr Endocrinol 4: 34–38. doi: 10.4274/jcrpe.473.

35. Farooqui AA, Horrocks LA, Farooqui T (2000a) Glycerophospholipids in brain: their metabolism, incorporation into membranes, functions, and involvement in neurological disorders. Chem Phys Lipids 106: 1–29. doi: 10.1016/s0009-3084(00)00128-6. PMID: 10878232.

36. Farooqui AA, Horrocks LA, Farooqui T (2000b) Deacylation and reacylation of neural membrane glycerophospholipids. J Mol Neurosci 14: 123–135. doi: 10.1385/jmn:14:3:123. PMID: 10984188.

37. Fedde KN, Blair L, Silverstein J, Coburn SP, Ryan LM, Weinstein RS, Waymire K, Narisawa S, Millán JL, MacGregor GR, Whyte MP (1999) Alkaline phosphatase knock-out mice recapitulate the metabolic and skeletal defects of infantile hypophosphatasia. J Bone Miner Res 14: 2015–2026. doi: 10.1359/jbmr.1999.14.12.2015.

38. Fitzner D, Bader JM, Penkert H, Bergner CG, Su M, Weil MT, Surma MA, Mann M, Klose C, Simons M (2020) Cell-typeand brain-region-resolved mouse brain lipidome. Cell Rep 32: 108132. doi: 10.1016/j.celrep.2020.108132. PMID: 32937123.

39. Fonta C, Négyessy L, Renaud L, Barone P (2004) Areal and subcellular localization of the ubiquitous alkaline phosphatase in the primate cerebral cortex: evidence for a role in neurotransmission. Cereb Cortex 14: 595–609. doi: 10.1093/cercor/bhh021. PMID: 15054075.

40. Fonta C, Negyessy L, Renaud L, Barone P (2005) Postnatal development of alkaline phosphatase activity correlates with the maturation of neurotransmission in the cerebral cortex. J Comp Neurol 486: 179–196. doi: 10.1002/cne.20524. PMID: 15844208.

41. Fonta C, Barone P, Rodriguez Martinez L, Négyessy L (2015) Rediscovering TNAP in the brain: a major role in regulating the function and development of the cerebral cortex. Subcell Biochem 76: 85–106. doi: 10.1007/978-94-017-7197-9_5.

42. Fraser D, Yendt ER, Christie FH (1955) Metabolic abnormalities in hypophosphatasia. Lancet 268: 286. doi: 10.1016/s0140-6736(55)90112-4.

43. Fraser D (1957) Hypophosphatasia. Am J Med 22: 730–746. doi: 10.1016/0002-9343(57)90124-9.

44. Friede RL (1966) A quantitative mapping of alkaline phosphatase in the brain of the rhesus monkey. J Neurochem 13: 197–203. doi: 10.1111/j.1471-4159.1966.tb07513.x. PMID: 4957290.

45. Fukazawa M, Tezuka J, Sasazuki M, Masumoto N, Baba H, Doi T, Tsutsumi Y, Mizuno Y, Mihara F, Nakayama H (2018) Infantile hypophosphatasia combined with vitamin B6-responsive seizures and reticular formation lesions on magnetic resonance imaging: A case report. Brain Dev 40: 140–144. doi: 10.1016/j.braindev.2017.07.015. PMID: 28802630.

46. Ghandour MS, Skoff RP (1988) Expression of galactocerebroside in developing normal and jimpy oligodendrocytes *in situ*. J Neurocytol 17: 485–498. doi: 10.1007/BF01189804. PMID: 3193128.

47. Goldstein DJ, Harris H (1981) Mammalian brain alkaline phosphatase: expression of liver/bone/kidney locus. Comparison of fetal and adult activities. J Neurochem 36: 53–57. doi: 10.1111/j.1471-4159.1981.tb02376.x. PMID: 7193242.

48. Green P, Glozman S, Kamensky B, Yavin E (1999) Developmental changes in rat brain membrane lipids and fatty acids. The preferential prenatal accumulation of docosahexaenoic acid. J Lipid Res 40: 960–966. PMID: 10224166.

49. Gregory JF, DeRatt BN, Rios-Avila L, Ralat M, Stacpoole PW (2016) Vitamin B6 nutritional status and cellular availability of pyridoxal 5’-phosphate govern the function of the transsulfuration pathway’s canonical reactions and hydrogen sulfide production via side reactions. Biochimie 126: 21–26. doi: 10.1016/j.biochi.2015.12.020. PMID: 26765812; PMCID: PMC4899128.

50. Grønbæk-Thygesen M, Hartmann-Petersen R (2024) Cellular and molecular mechanisms of aspartoacylase and its role in Canavan disease. Cell Biosci 14: 45. doi: 10.1186/s13578-024-01224-6. PMID: 38582917; PMCID: PMC10998430.

51. Guan ZZ, Wang YN, Xiao KQ, Hu PS, Liu JL (1999) Activity of phosphatidylethanolamine-N-methyltransferase in brain affected by Alzheimer’s disease. Neurochem Int 34: 41–47. doi: 10.1016/s0197-0186(98)00068-0. PMID: 10100195.

52. Hanada K (2003) Serine palmitoyltransferase, a key enzyme of sphingolipid metabolism. Biochim Biophys Acta 1632: 16–30. doi: 10.1016/s1388-1981(03)00059-3. PMID: 12782147.

53. Hanics J, Barna J, Xiao J, Millán JL, Fonta C, Négyessy L (2012) Ablation of TNAP function compromises myelination and synaptogenesis in the mouse brain. Cell Tissue Res 349: 459–471. doi: 10.1007/s00441-012-1455-z. PMID: 22696173; PMCID: PMC3415568.

54. Harayama T, Riezman H (2018) Understanding the diversity of membrane lipid composition. Nat Rev Mol Cell Biol 19: 281–296. doi: 10.1038/nrm.2017.138. PMID: 29410529.

55. Hargreaves KM, Clandinin MT (1989) Co-ordinate control of CDP-choline and phosphatidylethanolamine methyltransferase pathways for phosphatidylcholine biosynthesis occurs in response to change in diet fat. Biochim Biophys Acta 1001: 262–267. doi: 10.1016/0005-2760(89)90109-4. PMID: 2917151.

56. Hilbig R, Rösner H, Merz G, Segler-Stahl K, Rahmann H (1982) Developmental profiles of gangliosides in mouse and rat cerebral cortex. Wilehm Roux Arch Dev Biol 191: 281–284. doi: 10.1007/BF00848417. PMID: 28305060.

57. Hofmann C, Liese J, Schwarz T, Kunzmann S, Wirbelauer J, Nowak J, Hamann J, Girschick H, Graser S, Dietz K, Zeck S, Jakob F, Mentrup B (2013) Compound heterozygosity of two functional null mutations in the ALPL gene associated with deleterious neurological outcome in an infant with hypophosphatasia. Bone 55: 150–157. doi: 10.1016/j.bone.2013.02.017.

58. Hoffman DR, Haning JA, Cornatzer WE (1981) Microsomal phosphatidylethanolamine methyltransferase: inhibition by S-adenosylhomocysteine. Lipids 16: 561–567. doi: 10.1007/BF02534900. PMID: 7278517.

59. Hoover-Fong J, Sobreira N, Jurgens J, Modaff P, Blout C, Moser A, Kim OH, Cho TJ, Cho SY, Kim SJ, Jin DK, Kitoh H, Park WY, Ling H, Hetrick KN, Doheny KF, Valle D, Pauli RM (2014) Mutations in *PCYT1A*, encoding a key regulator of phosphatidylcholine metabolism, cause spondylometaphyseal dysplasia with cone-rod dystrophy. Am J Hum Genet 94: 105–112. doi: 10.1016/j.ajhg.2013.11.018. PMID: 24387990; PMCID: PMC3882727.

60. Horrocks LA (1968) Composition of mouse brain myelin during development. J Neurochem 15: 483–488. doi: 10.1111/j.1471-4159.1968.tb08945.x. PMID: 5668048.

61. Isa Y, Tsuge H, Hayakawa T (2006a) Effect of vitamin B6 deficiency on S-adenosylhomocysteine hydrolase activity as a target point for methionine metabolic regulation. J Nutr Sci Vitaminol (Tokyo) 52: 302–306. doi: 10.3177/jnsv.52.302.

62. Isa Y, Mishima T, Tsuge H, Hayakawa T (2006b) Increase in S-adenosylhomocysteine content and its effect on the S-adenosylhomocysteine hydrolase activity under transient high plasma homocysteine levels in rats. J Nutr Sci Vitaminol (Tokyo) 52: 479–482. doi: 10.3177/jnsv.52.479.

63. Jain M, Ngoy S, Sheth SA, Swanson RA, Rhee EP, Liao R, Clish CB, Mootha VK, Nilsson R (2014) A systematic survey of lipids across mouse tissues. Am J Physiol Endocrinol Metab 306: E854–E868. doi: 10.1152/ajpendo.00371.2013. PMID: 24518676; PMCID: PMC3989739.

64. Johnson PI, Blusztajn JK (1998) Sexually dimorphic activation of liver and brain phosphatidylethanolamine Nmethyltransferase by dietary choline deficiency. Neurochem Res 23: 583–587. doi: 10.1023/a:1022470301550. PMID: 9566595.

65. Kennedy EP, Weiss SB (1956) The function of cytidine coenzymes in the biosynthesis of phospholipides. J Biol Chem 222: 193–214. PMID: 13366993.

66. Kermer V, Ritter M, Albuquerque B, Leib C, Stanke M, Zimmermann H (2010) Knockdown of tissue nonspecific alkaline phosphatase impairs neural stem cell proliferation and differentiation. Neurosci Lett 485: 208–211. doi: 10.1016/j.neulet.2010.09.013. PMID: 20849921.

67. Kishimoto Y, Agranoff BW, Radin NS, Burton RM (1969) Comparison of the fatty acids of lipids of subcellular brain fractions. J Neurochem 16: 397–404. doi: 10.1111/j.1471-4159.1969.tb10380.x. PMID: 5802640.

68. Kitagawa E, Yamamoto T, Yamamoto K, Nakagawa T, Hayakawa T (2015) Accumulation of lipid in rat liver was induced by vitamin B₆ deficiency and was ameliorated by supplemental phosphatidylcholine in the diet. Biosci Biotechnol Biochem 79: 1320–1326. doi: 10.1080/09168451.2015.1020755. PMID: 25775923.

69. Klöckner C, Fernández-Murray JP, Tavasoli M, Sticht H, Stoltenburg-Didinger G, Scholle LM, Bakhtiari S, Kruer MC, Darvish H, Firouzabadi SG, Pagnozzi A, Shukla A, Girisha KM, Narayanan DL, Kaur P, Maroofian R, Zaki MS, Noureldeen MM, Merkenschlager A, Gburek-Augustat J, Cali E, Banu S, Nahar K, Efthymiou S, Houlden H, Jamra RA, Williams J, McMaster CR, Platzer K (2022) Bi-allelic variants in *CHKA* cause a neurodevelopmental disorder with epilepsy and microcephaly. Brain 145: 1916–1923. doi: 10.1093/brain/awac074. PMID: 35202461; PMCID: PMC9630884.

70. Kloor D, Osswald H (2004) S-Adenosylhomocysteine hydrolase as a target for intracellular adenosine action. Trends Pharmacol Sci 25: 294–297. doi: 10.1016/j.tips.2004.04.004.

71. Kurtz DJ, Levy H, Kanfer JN (1972) Cerebral lipids and amino acids in the vitamin B_6_-deficient suckling rat. J Nutr 102: 291–298. doi: 10.1093/jn/102.2.291.

72. Kurtz DJ, Kanfer JN (1973) Composition of myelin lipids and synthesis of 3-ketodihydrosphingosine in the vitamin B_6_-deficient developing rat. J Neurochem 20: 963–968. doi: 10.1111/j.1471-4159.1973.tb00066.x. PMID: 4697895.

73. Lakher M, Wurtman RJ (1987) *In vivo* synthesis of phosphatidylcholine in rat brain via the phospholipid methylation pathway. Brain Res 419: 131–140. doi: 10.1016/0006-8993(87)90576-2. PMID: 3676720.

74. Langer D, Ikehara Y, Takebayashi H, Hawkes R, Zimmermann H (2007) The ectonucleotidases alkaline phosphatase and nucleoside triphosphate diphosphohydrolase 2 are associated with subsets of progenitor cell populations in the mouse embryonic, postnatal and adult neurogenic zones. Neuroscience 150: 863–879. doi: 10.1016/j.neuroscience.2007.07.064. PMID: 18031938.

75. Langer D, Hammer K, Koszalka P, Schrader J, Robson S, Zimmermann H (2008) Distribution of ectonucleotidases in the rodent brain revisited. Cell Tissue Res 334: 199–217. doi: 10.1007/s00441-008-0681-x. PMID: 18843508.

76. Lauwers E, Goodchild R, Verstreken P (2016) Membrane lipids in presynaptic function and disease. Neuron 90: 11–25. doi: 10.1016/j.neuron.2016.02.033. PMID: 27054615.

77. Li Z, Agellon LB, Allen TM, Umeda M, Jewell L, Mason A, Vance DE (2006) The ratio of phosphatidylcholine to phosphatidylethanolamine influences membrane integrity and steatohepatitis. Cell Metab 3: 321–331. doi: 10.1016/j.cmet.2006.03.007. PMID: 16679290.

78. Liebisch G, Fahy E, Aoki J, Dennis EA, Durand T, Ejsing CS, Fedorova M, Feussner I, Griffiths WJ, Köfeler H, Merrill AH Jr, Murphy RC, O’Donnell VB, Oskolkova O, Subramaniam S, Wakelam MJO, Spener F (2020) Update on LIPID MAPS classification, nomenclature, and shorthand notation for MS-derived lipid structures. J Lipid Res 61: 1539–1555. doi: 10.1194/jlr.S120001025. PMID: 33037133; PMCID: PMC7707175.

79. Lillington JM, Trafford DJ, Makin HL (1981) A rapid and simple method for the esterification of fatty acids and steroid carboxylic acids prior to gas-liquid chromatography. Clin Chim Acta 111: 91–98. doi: 10.1016/0009-8981(81)90425-3. PMID: 7226543.

80. Loenen WA (2006) S-adenosylmethionine: jack of all trades and master of everything? Biochem Soc Trans 34: 330–333. doi: 10.1042/BST20060330. PMID: 16545107.

81. Loo G, Smith JT (1986) Effect of pyridoxine deficiency on phospholipid methylation in rat liver microsomes. Lipids 21: 409–412. doi: 10.1007/BF02534937. PMID: 3736351.

82. Marcucci H, Paoletti L, Jackowski S, Banchio C (2010) Phosphatidylcholine biosynthesis during neuronal differentiation and its role in cell fate determination. J Biol Chem 285: 25382–25393. doi: 10.1074/jbc.M110.139477. PMID: 20525991; PMCID: PMC2919101.

83. Martinez M, Cuskelly GJ, Williamson J, Toth JP, Gregory JF 3rd (2000) Vitamin B-6 deficiency in rats reduces hepatic serine hydroxymethyltransferase and cystathionine beta-synthase activities and rates of in vivo protein turnover, homocysteine remethylation and transsulfuration. J Nutr 130: 1115–1123. doi: 10.1093/jn/130.5.1115.

84. Millán JL (2006a). Mammalian alkaline phosphatases. From biology to applications in medicine and biotechnology. Wiley-VCH Verlag GmbH & Co, Weinheim, Germany.

85. Millán JL (2006b) Alkaline phosphatases: structure, substrate specificity and functional relatedness to other members of a large superfamily of enzymes. Purinergic Signal 2: 335–341. doi: 10.1007/s11302-005-5435-6. PMID: 18404473; PMCID: PMC2254479.

86. Montaner A, da Silva Santana TT, Schroeder T, Einicker-Lamas M, Girardini J, Costa MR, Banchio C (2018) Specific Phospholipids Regulate the Acquisition of Neuronal and Astroglial Identities in Post-Mitotic Cells. Sci Rep 8: 460. doi: 10.1038/s41598-017-18700-4. PMID: 29323239; PMCID: PMC5765016.

87. Mori S, Nagano M (1985) Electron microscopic cytochemistry of alkaline phosphatase in neurons of rats. Arch Histol Jpn 48: 389–397. doi: 10.1679/aohc.48.389. PMID: 4084005.

88. Mornet E, Yvard A, Taillandier A, Fauvert D, Simon-Bouy B (2011) A molecular-based estimation of the prevalence of hypophosphatasia in the European population. Ann Hum Genet 75: 439–445. doi: 10.1111/j.1469-1809.2011.00642.x. PMID: 21488855.

89. Morré DM, Kirksey A, Das GD (1978) Effects of vitamin B-6 deficiency on the developing central nervous system of the rat. Myelination. J Nutr 108: 1260–1265. doi: 10.1093/jn/108.8.1260.

90. Motulsky H (1995) Intuitive Biostatistics. Oxford University Press.

91. Mitsuhashi S, Ohkuma A, Talim B, Karahashi M, Koumura T, Aoyama C, Kurihara M, Quinlivan R, Sewry C, Mitsuhashi H, Goto K, Koksal B, Kale G, Ikeda K, Taguchi R, Noguchi S, Hayashi YK, Nonaka I, Sher RB, Sugimoto H, Nakagawa Y, Cox GA, Topaloglu H, Nishino I (2011) A congenital muscular dystrophy with mitochondrial structural abnormalities caused by defective *de novo* phosphatidylcholine biosynthesis. Am J Hum Genet 88: 845–851. doi: 10.1016/j.ajhg.2011.05.010. PMID: 21665002; PMCID: PMC3113344.

92. Naffaa V, Magny R, Regazzetti A, Van Steenwinckel J, Gressens P, Laprévote O, Auzeil N, Schang AL (2022) Shift in phospholipid and fatty acid contents accompanies brain myelination. Biochimie 203: 20–31. doi: 10.1016/j.biochi.2022.08.010. PMID: 36055603.

93. Namboodiri AM, Peethambaran A, Mathew R, Sambhu PA, Hershfield J, Moffett JR, Madhavarao CN (2006) Canavan disease and the role of N-acetylaspartate in myelin synthesis. Mol Cell Endocrinol 252: 216–223. doi: 10.1016/j.mce.2006.03.016. PMID: 16647192.

94. Narisawa S, Hasegawa H, Watanabe K, Millán JL (1994) Stage-specific expression of alkaline phosphatase during neural development in the mouse. Dev Dyn 201: 227–235. doi: 10.1002/aja.1002010306. PMID: 7533563.

95. Narisawa S, Fröhlander N, Millán JL (1997) Inactivation of two mouse alkaline phosphatase genes and establishment of a model of infantile hypophosphatasia. Dev Dyn 208: 432–446. doi: 10.1002/(SICI)10970177(199703)208:3<432::AID-AJA13>3.0.CO;2-1. PMID: 9056646.

96. Négyessy L, Xiao J, Kántor O, Kovács GG, Palkovits M, Dóczi TP, Renaud L, Baksa G, Glasz T, Ashaber M, Barone P, Fonta C (2011) Layer-specific activity of tissue non-specific alkaline phosphatase in the human neocortex. Neuroscience 172: 406–418. doi: 10.1016/j.neuroscience.2010.10.049. PMID: 20977932.

97. Nguyen TT, Hayakawa T, Tsuge H (2001) Effect of vitamin B6 deficiency on the synthesis and accumulation of S-adenosylhomocysteine and S-adenosylmethionine in rat tissues. J Nutr Sci Vitaminol (Tokyo) 47: 188–194. doi: 10.3177/jnsv.47.188. PMID: 11575573.

98. Nitsch RM, Blusztajn JK, Pittas AG, Slack BE, Growdon JH, Wurtman RJ (1992) Evidence for a membrane defect in Alzheimer disease brain. Proc Natl Acad Sci USA 89: 1671–1675. doi: 10.1073/pnas.89.5.1671. PMID: 1311847; PMCID: PMC48514.

99. Norton WT, Poduslo SE (1973) Myelination in rat brain: changes in myelin composition during brain maturation. J Neurochem 21: 759–773. doi: 10.1111/j.1471-4159.1973.tb07520.x. PMID: 4754856.

100. O’Brien JS, Sampson EL (1965a) Lipid composition of the normal human brain: gray matter, white matter, and myelin. J Lipid Res 6: 537–544. PMID: 5865382.

101. O’Brien JS, Sampson EL (1965b) Fatty acid and fatty aldehyde composition of the major brain lipids in normal human gray matter, white matter, and myelin. J Lipid Res 6: 545–551. PMID: 5865383.

102. O’Donnell VB (2022) New appreciation for an old pathway: the Lands Cycle moves into new arenas in health and disease. Biochem Soc Trans 50: 1–11. doi: 10.1042/BST20210579. PMID: 35225335; PMCID: PMC9022965.

103. Oosterink JE, Naninck EF, Korosi A, Lucassen PJ, van Goudoever JB, Schierbeek H (2015) Accurate measurement of the essential micronutrients methionine, homocysteine, vitamins B6, B12, B9 and their metabolites in plasma, brain and maternal milk of mice using LC/MS ion trap analysis. J Chromatogr B 998-999: 106–113. doi: 10.1016/j.jchromb.2015.07.008. PMID: 26197436.

104. Osetrova M, Tkachev A, Mair W, Guijarro Larraz P, Efimova O, Kurochkin I, Stekolshchikova E, Anikanov N, Foo JC, Cazenave-Gassiot A, Mitina A, Ogurtsova P, Guo S, Potashnikova DM, Gulin AA, Vasin AA, Sarycheva A, Vladimirov G, Fedorova M, Kostyukevich Y, Nikolaev E, Wenk MR, Khrameeva EE, Khaitovich P (2024) Lipidome atlas of the adult human brain. Nat Commun 15: 4455. doi: 10.1038/s41467-024-48734-y. PMID: 38796479; PMCID: PMC11127996.

105. Paoletti L, Domizi P, Marcucci H, Montaner A, Krapf D, Salvador G, Banchio C (2016) Lysophosphatidylcholine drives neuroblast cell fate. Mol Neurobiol 53: 6316–6331. doi: 10.1007/s12035-015-9528-0. PMID: 26567110.

106. Percudani R, Peracchi A (2009) The B6 database: a tool for the description and classification of vitamin B6-dependent enzymatic activities and of the corresponding protein families. BMC bioinformatics 10: 273. doi: 10.1186/1471-2105-10-273. PMID: 19723314; PMCID: PMC2748086.

107. Phillips GR, Hancock SE, Jenner AM, McLean C, Newell KA, Mitchell TW (2022) Phospholipid profiles are selectively altered in the putamen and white frontal cortex of Huntington’s disease. Nutrients 14: 2086. doi: 10.3390/nu14102086. PMID: 35631226; PMCID: PMC9143248.

108. Phillis JW, Edstrom JP, Kostopoulos GK, Kirkpatrick JR (1979) Effects of adenosine and adenine nucleotides on synaptic transmission in the cerebral cortex. Can J Physiol Pharmacol 57: 1289–1312. doi: 10.1139/y79-194.

109. Phoenix KN, Marozsan A, Liu-Chen S, Zhang L, Mogha A, Wong Y, Brownstein CA, Padidela R (2025) Asfotase alfa restores PLP-dependent GABA, cystathionine, and amino acid metabolism in a mouse model of hypophosphatasia. Neurosci Res 218: 104940. doi: 10.1016/j.neures.2025.104940.

110. Posse de Chaves E, Vance DE, Campenot RB, Vance JE (1995a). Axonal synthesis of phosphatidylcholine is required for normal axonal growth in rat sympathetic neurons. J Cell Biol 128: 913–918. doi: 10.1083/jcb.128.5.913. PMID: 7876314; PMCID: PMC2120403.

111. Posse de Chaves E, Vance DE, Campenot RB, Vance JE (1995b) Alkylphosphocholines inhibit choline uptake and phosphatidylcholine biosynthesis in rat sympathetic neurons and impair axonal extension. Biochem J 312: 411–417. doi: 10.1042/bj3120411. PMID: 8526849; PMCID: PMC1136277.

112. Pynn CJ, Henderson NG, Clark H, Koster G, Bernhard W, Postle AD (2011) Specificity and rate of human and mouse liver and plasma phosphatidylcholine synthesis analyzed in vivo. J Lipid Res 52: 399–407. doi: 10.1194/jl-r.D011916. PMID: 21068006; PMCID: PMC3023562.

113. Rathbun JC (1948) Hypophosphatasia; a new developmental anomaly. Am J Dis Child 75: 822–831. doi: 10.1001/archpedi.1948.02030020840003.

114. Reddington M, Pusch R (1983) Adenosine metabolism in a rat hippocampal slice preparation: incorporation into S-adenosylhomocysteine. J Neurochem 40: 285–290. doi: 10.1111/j.1471-4159.1983.tb12684.x.

115. Rickman OJ, Baple EL, Crosby AH (2020) Lipid metabolic pathways converge in motor neuron degenerative diseases. Brain 143: 1073–1087. doi: 10.1093/brain/awz382. PMID: 31848577; PMCID: PMC7174042.

116. Russell RG (1965) excretion of inorganic pyrophosphate in hypophosphatasia. Lancet 2 (7410): 461–464. doi: 10.1016/s0140-6736(65)91422-4.

117. Salles JP (2020) Hypophosphatasia: biological and clinical aspects, avenues for therapy. Clin Biochem Rev 41: 13–27. doi: 10.33176/AACB-19-00031. PMID: 32158059; PMCID: PMC7043401.

118. Schmitt S, Castelvetri LC, Simons M (2015) Metabolism and functions of lipids in myelin. Biochim Biophys Acta 1851: 999–1005. doi: 10.1016/j.bbalip.2014.12.016. PMID: 25542507.

119. Schonbach J, Hu KH, Friede RL (1968) Cellular and chemical changes during myelination: histologic, autoradiographic, histochemical and biochemical data on myelination in the pyramidal tract and corpus callosum of rat. J Comp Neurol 134: 21–36. doi: 10.1002/cne.901340104. PMID: 4181214.

120. Sebastián-Serrano Á, Engel T, de Diego-García L, Olivos-Oré LA, Arribas-Blázquez M, Martínez-Frailes C, Pérez-Díaz C, Millán JL, Artalejo AR, Miras-Portugal MT, Henshall DC, Díaz-Hernández M (2016) Neurodevelopmental alterations and seizures developed by mouse model of infantile hypophosphatasia are associated with purinergic signalling deregulation. Hum Mol Genet 25: 4143–4156. doi: 10.1093/hmg/ddw248. PMID: 27466191; PMCID: PMC5291194.

121. She QB, Hayakawa T, Tsuge H (1995) Alteration in the phosphatidylcholine biosynthesis of rat liver microsomes caused by vitamin B6 deficiency. Biosci Biotechnol Biochem 59: 163–167. doi: 10.1271/bbb.59.163. PMID: 7766014.

122. Shiota C, Miura M, Mikoshiba K (1989) Developmental profile and differential localization of mRNAs of myelin proteins (MBP and PLP) in oligodendrocytes in the brain and in culture. Brain Res Dev Brain Res 45: 83–94. doi: 10.1016/0165-3806(89)90010-2. PMID: 2465104.

123. Sia C, Wapnir R, Sokal M, Harper RG, Intizar S, Manhasset F (1975) Effects of pyridoxine on neonatal hypophosphatasia. Pediatr Res 9: 355.

124. Sjöstedt E, Zhong W, Fagerberg L, Karlsson M, Mitsios N, Adori C, Oksvold P, Edfors F, Limiszewska A, Hikmet F, Huang J, Du Y, Lin L, Dong Z, Yang L, Liu X, Jiang H, Xu X, Wang J, Yang H, Bolund L, Mardinoglu A, Zhang C, von Feilitzen K, Lindskog C, Pontén F, Luo Y, Hökfelt T, Uhlén M, Mulder J (2020) An atlas of the protein-coding genes in the human, pig, and mouse brain. Science 367: eaay5947. doi: 10.1126/science.aay5947. PMID: 32139519.

125. Stevens B, Porta S, Haak LL, Gallo V, Fields RD (2002) Adenosine: a neuron-glial transmitter promoting myelination in the CNS in response to action potentials. Neuron 36: 855–868. doi: 10.1016/s0896-6273(02)01067-x.

126. Sugimura K, Mizutani A (1979) Histochemical and cytochemical studies of alkaline phosphatase activity in the synapses of rat brain. Histochemistry 61: 123–129. doi: 10.1007/BF00496524. PMID: 457450.

127. Suzuki K, Okada M (1982) Alterations of phospholipid and triglyceride metabolism in fatty liver caused by pyridoxine deficiency in rats. J Nutr Sci Vitaminol (Tokyo) 28: 377–390. doi: 10.3177/jnsv.28.377. PMID: 7175577.

128. Taketani T, Onigata K, Kobayashi H, Mushimoto Y, Fukuda S, Yamaguchi S (2014) Clinical and genetic aspects of hypophosphatasia in Japanese patients. Arch Dis Child 99: 211–215. doi: 10.1136/archdischild-2013-305037.

129. Tashiro S, Sudou K, Imoh A, Koide M, Akazawa Y (1983) Phosphatidylethanolamine methyltransferase activity in developing, demyelinating, and diabetic mouse brain. Tohoku J Exp Med 141, Suppl: 485–490. doi: 10.1620/tjem.141.suppl_485. PMID: 6680524.

130. Tavasoli M, Lahire S, Reid T, Brodovsky M, McMaster CR (2020) Genetic diseases of the Kennedy pathways for membrane synthesis. J Biol Chem 295: 17877–17886. doi: 10.1074/jbc.REV120.013529. PMID: 33454021; PMCID: PMC7762932.

131. Valentine WJ, Shimizu T, Shindou H (2023) Lysophospholipid acyltransferases orchestrate the compositional diversity of phospholipids. Biochimie 215: 24–33. doi: 10.1016/j.biochi.2023.08.012. PMID: 37611890.

132. van der Stoep N, Onkenhout W, Prins S, Struys E, Jakobs C, Peeters-Scholte C (2014) Comments on ’Infantile hypophosphatasia without bone deformities presenting with severe pyridoxine-resistant seizures’ in Molecular Genetics and Metabolism’ 2014 Mar;111(3):404–7 by M.G. de Roo, N.G. Abeling, C.B. Majoie, A.M. Bosch, J.H. Koelman, J.M. Cobben, M. Duran, B.T. Poll-The. Mol Genet Metab Rep 1: 220-222. doi: 10.1016/j.ymgmr.2014.04.005. PMID: 27896090; PMCID: PMC5121321.

133. van der Veen JN, Kennelly JP, Wan S, Vance JE, Vance DE, Jacobs RL (2017) The critical role of phosphatidylcholine and phosphatidylethanolamine metabolism in health and disease. Biochim Biophys Acta Biomemb 1859 : 1558–1572. doi: 10.1016/j.bbamem.2017.04.006. PMID: 28411170.

134. Vance JE, Pan D, Campenot RB, Bussière M, Vance DE (1994) Evidence that the major membrane lipids, except cholesterol, are made in axons of cultured rat sympathetic neurons. J Neurochem 62: 329–337. doi: 10.1046/j.1471-4159.1994.62010329.x. PMID: 8263532.

135. Walkey CJ, Donohue LR, Bronson R, Agellon LB, Vance DE (1997) Disruption of the murine gene encoding phosphatidylethanolamine N-methyltransferase. Proc Natl Acad Sci USA 94: 12880–12885. doi: 10.1073/pnas.94.24.12880. PMID: 9371769; PMCID: PMC24232.

136. Wallace TC, Blusztajn JK, Caudill MA, Klatt KC, Zeisel SH (2020) Choline: The neurocognitive essential nutrient of interest to obstetricians and gynecologists. J Diet Suppl 17: 733–752. doi: 10.1080/19390211.2019.1639875. PMID: 31385730.

137. Waymire KG, Mahuren JD, Jaje JM, Guilarte TR, Coburn SP, MacGregor GR (1995) Mice lacking tissue nonspecific alkaline phosphatase die from seizures due to defective metabolism of vitamin B-6. Nat Genet 11: 45–51. doi: 10.1038/ng0995-45. PMID: 7550313.

138. Wells MA, Dittmer JC (1967) A comprehensive study of the postnatal changes in the concentration of the lipids of developing rat brain. Biochemistry 6: 3169–3175. doi: 10.1021/bi00862a026. PMID: 6056982.

139. Whyte MP, Mahuren JD, Vrabel LA, Coburn SP (1985) Markedly increased circulating pyridoxal-5’-phosphate levels in hypophosphatasia. Alkaline phosphatase acts in vitamin B6 metabolism. J Clin Invest 76: 752–756. doi: 10.1172/JCI112031. PMID: 4031070; PMCID: PMC423894.

140. Whyte MP (2010) Physiological role of alkaline phosphatase explored in hypophosphatasia. Ann N Y Acad Sci 1192: 190–200. doi: 10.1111/j.1749-6632.2010.05387.x.

141. Whyte MP, Rockman-Greenberg C, Ozono K, Riese R, Moseley S, Melian A, Thompson DD, Bishop N, Hofmann C (2016) Asfotase Alfa Treatment Improves Survival for Perinatal and Infantile Hypophosphatasia. J Clin Endocrinol Metab 101: 334–342. doi: 10.1210/jc.2015-3462. PMID: 26529632.

142. Williamson B, Coniglio JG (1971) The effects of pyridoxine deficiency and of caloric restriction on lipids in the developing rat brain. J Neurochem 18: 267–276. doi: 10.1111/j.1471-4159.1971.tb00565.x. PMID: 5550092.

143. Wood PL (2012) Lipidomics of Alzheimer’s disease: current status. Alzheimers Res Ther 4: 5. doi: 10.1186/alzrt103. PMID: 22293144; PMCID: PMC3471525.

144. Xicoy H, Wieringa B, Martens GJM (2019) The role of lipids in Parkinson’s disease. Cells 8: 27. doi: 10.3390/cells8010027. PMID: 30621069; PMCID: PMC6356353.

145. Yoon JH, Seo Y, Jo YS, Lee S, Cho E, Cazenave-Gassiot A, Shin YS, Moon MH, An HJ, Wenk MR, Suh PG (2022) Brain lipidomics: From functional landscape to clinical significance. Sci Adv 8: eadc9317. doi: 10.1126/sciadv.adc9317. PMID: 36112688; PMCID: PMC9481132.

146. Zhu X, Mar MH, Song J, Zeisel SH (2004) Deletion of the Pemt gene increases progenitor cell mitosis, DNA and protein methylation and decreases calretinin expression in embryonic day 17 mouse hippocampus. Brain Res Dev Brain Res 149: 121–129. doi: 10.1016/j.devbrainres.2004.01.004. PMID: 15063092.

